# Predicting clonal somatic mutations from tumour spatial transcriptomics data using a graph convolutional neural network

**DOI:** 10.64898/2026.07.08.737173

**Authors:** Tom Luijts, Sofie Hoogstoel, Song Wen, Eden Pappaert, Ellen De Meester, Filip Van Nieuwerburgh, Evelien Van Hamme, Sofie De Schepper, Wouter Willaert, Anne Vral, Isabelle Hoorens, Jimmy Van den Eynden

**Affiliations:** Department of Human Structure and Repair, Ghent University, Ghent, Belgium; Cancer Research Institute Ghent, Ghent, Belgium; NXTGNT, Ghent University, Ghent, Belgium; VIB Spatial Catalyst, VIB Technologies, VIB, Ghent, Belgium; Department of Dermatology, University Hospital Ghent; Belgium; Department of Gastrointestinal Surgery, Ghent University Hospital, Ghent, Belgium

## Abstract

Spatial transcriptomics (ST) has revolutionized our understanding of tumour biology but inherently lacks information on the upstream somatic driver mutations. We developed a spatially-aware graph convolutional neural network approach (SPOT-Mut) that infers mutated (sub)clones directly from ST data. SPOT-Mut was trained on virtual ST slides with clones simulated from a large collection of existing RNA and matched DNA sequencing data. SPOT-Mut was first evaluated for predicting mutations in *TP53*, the most frequently mutated gene in human cancer. Precision and recall values exceeded 95% in most analysed cancer types and the model was validated on ST and matched DNA sequencing data from serial slices obtained from cutaneous SCC samples. SPOT-Mut was then trained on 62 frequently mutated driver genes, resulting in highly performing somatic mutation prediction models for 40 driver genes, including *KRAS*, *KEAP1*, *CTNNB1* and *APC*. With the increasing availability of ST data and upcoming atlases, SPOT-Mut potentiates the localization of (sub)clonal alterations in *TP53* and other common driver mutations.

## Introduction

Spatial transcriptomics (ST) has successfully integrated the microscopic and transcriptomic information layers, providing a more holistic understanding of tissue properties. Systematic analyses of whole-transcriptome ST data derived from sequence-based technologies (e.g., *Visium* (*HD*), *Slide-Seq (V2)* or *Stereo-Seq*) provided new insights in the tumour and its microenvironment including paracrine signaling, tumour-immune interactions and cellular subpopulations and niches (1–3).

While an increasing amount of these tumour ST data are being made publicly available, resulting in secondary discoveries (4–6), they inherently lack key information on the upstream genomic events that are driving the tumour. Copy number alterations (CNAs) can still be inferred from ST data by combining genomic information with gene expression changes (7,8). However, this is not possible for single nucleotide variants (SNVs).

It was previously suggested that mutations in canonical cancer driver genes such as *TP53* and Ras pathway oncogenes (*KRAS, HRAS and NRAS*) can be predicted using machine learning models applied on bulk transcriptomics data (9,10). However, to our knowledge, such prediction models have not been created on ST data. Given the (sub)clonal nature of cells carrying driver mutations, neighboring spots on an ST slide have a higher probability to carry the same mutation and we hypothesized that the incorporation of this spatial information vastly increases the prediction accuracy.

We first focused on the p53 protein, encoded by the *TP53* gene. *TP53* is located on chromosome 17p13.1 and is mutated in about half of all tumours and one of the best-studied tumour suppressor genes (11–13). It’s a transcription factor central in the regulation of cellular response to DNA damage and stress, playing a crucial role in regulating cell cycle arrest, DNA damage repair and apoptosis (14–16). The activity of p53 is tightly regulated, primarily through its negative regulator Mdm2, an E3 ubiquitin ligase that targets p53 for degradation and controls its stability under normal conditions (17–19). Functional p53 is crucial for genomic stability (20,21) and because of these versatility of functions it is considered the guardian of the genome (22). *TP53* mutations are often correlated with poor prognosis and aggressive tumour characteristics (23–25). Somatic mutations in related genes have been shown to result in downstream phenotypic consequences that resemble those caused by mutated *TP53*. Such *TP53*-like alterations are referred to as phenocopies and include alterations in *MDM2*, *MDM4*, *PPM1D* or *USP28* (10,26–31).

To predict *TP53* and other somatic mutations in ST data, we developed SPOT-Mut (Spatial Prediction Of Tumor Mutations), a graph convolutional neural network (GCNN) that inherently models the spatial relation between neighboring spots. We demonstrate that this approach can accurately predict simulated mutated (sub)clones for 40 common driver genes and has superior performance compared to non-spatial approaches. Furthermore, we show that SPOT-Mut generalizes across cancer types and independent datasets and captures biologically meaningful transcriptional programs. Together, these results highlight the potential of SPOT-Mut as a methodological framework for spatially resolving somatic mutations from transcriptomic data in the absence of direct spatial genomic profiles.

## Results

### SPOT-Mut is trained on simulated spatial transcriptomics slides

Training a deep learning model to predict somatic mutations from ST data would require the availability of matched ST and spatially resolved genomics reference data. As spatial genomics technologies are still in their infancy (7,32), such training data are not (yet) available. To overcome this problem, we opted to train the model on virtual ST slides, generated from matched bulk DNA and RNA sequencing data. This approach was first evaluated for *TP53*, the most frequently mutated gene in human cancer. Bulk RNA-seq data were downloaded from The Cancer Genome Atlas (TCGA) along with the mutation status of the *TP53* gene (n = 6563; samples with both expression and mutation data). Subjects were randomly grouped in a training (70%) and test cohort (30%). After simulating 7 *TP53* subclones on an *in-silico* generated virtual ST slide, cancer type-specific TCGA training data were randomly assigned to ST spots while considering *TP53* mutation status. Finally, the grid of spots was converted to an undirected graph by connecting each spot to its neighbouring spots to form edges (connections) and nodes (spots). These graphs were used to train a GCNN followed by a multilayer perceptron (MLP) for binary classification of each node (Fig. 1).

**Fig. 1.**
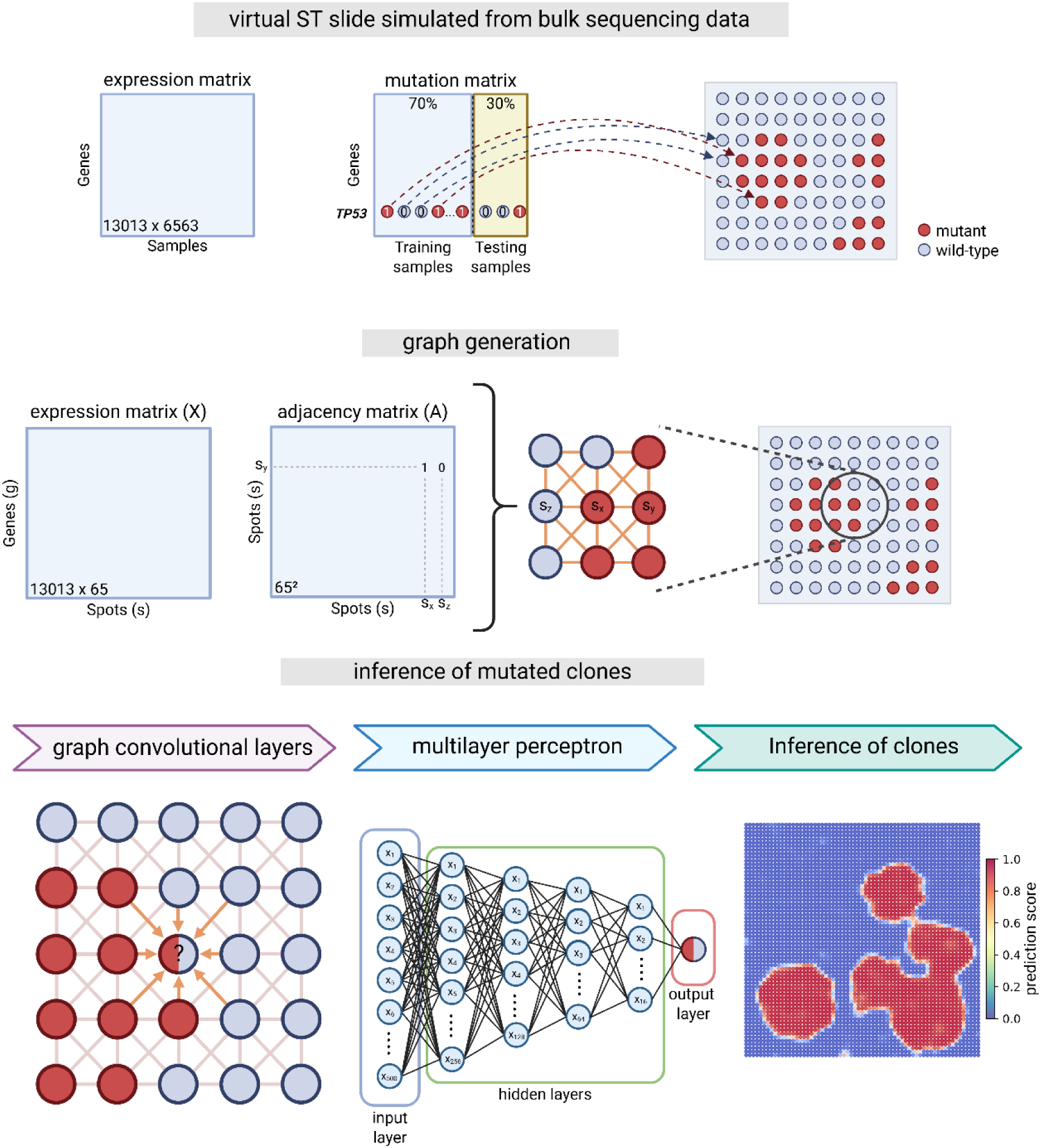
Overview of the *TP53* SPOT-Mut training process on simulated spatial transcriptomics slides Input data consist of a gene expression matrix containing FPKM normalized count values and a binary mutation matrix containing the mutation status of each gene in each sample. Data were derived from the cancer genome atlas (TCGA) and were randomly split in training (70%) and testing (30%) samples. Mutation status was used to assign each sample to a random spatial location on a virtual spatial transcriptomics (ST) slide containing simulated *TP53* clones. A graph can then be generated which connects each spot on the ST slide to its 8 neighbouring spots, resulting in an adjacency matrix (A) and a gene expression matrix (X). SPOT-Mut passes both A and X through two graph convolutional layers, along the connections defined in matrix A. The output of the final graph convolutional layer is passed through a feed-forward multilayer perceptron with four hidden layers, resulting in the prediction of the spatial location of *TP53* mutant spots.

### SPOT-Mut accurately predicts single and double hit *TP53* mutations in different cancer types

The prediction accuracy of the trained model was determined using the test data (n = 1979). For each individual cancer type, as well as the mixed, pan-cancer dataset, 100 ST test slides were simulated and fed to the model for binary prediction of each spot (*TP53* mutated or wild-type). The model accurately predicted *TP53* mutation status for the pan-cancer datasets (precision 0.97; recall 0.97; Fig. 2a-b) as well as the 12 individually analysed TCGA cancer types (median precision 0.97; median recall 0.96). Recall was lowest for SKCM (0.86) while STAD had the lowest precision value (0.89; Fig. 2b).

**Fig. 2.**
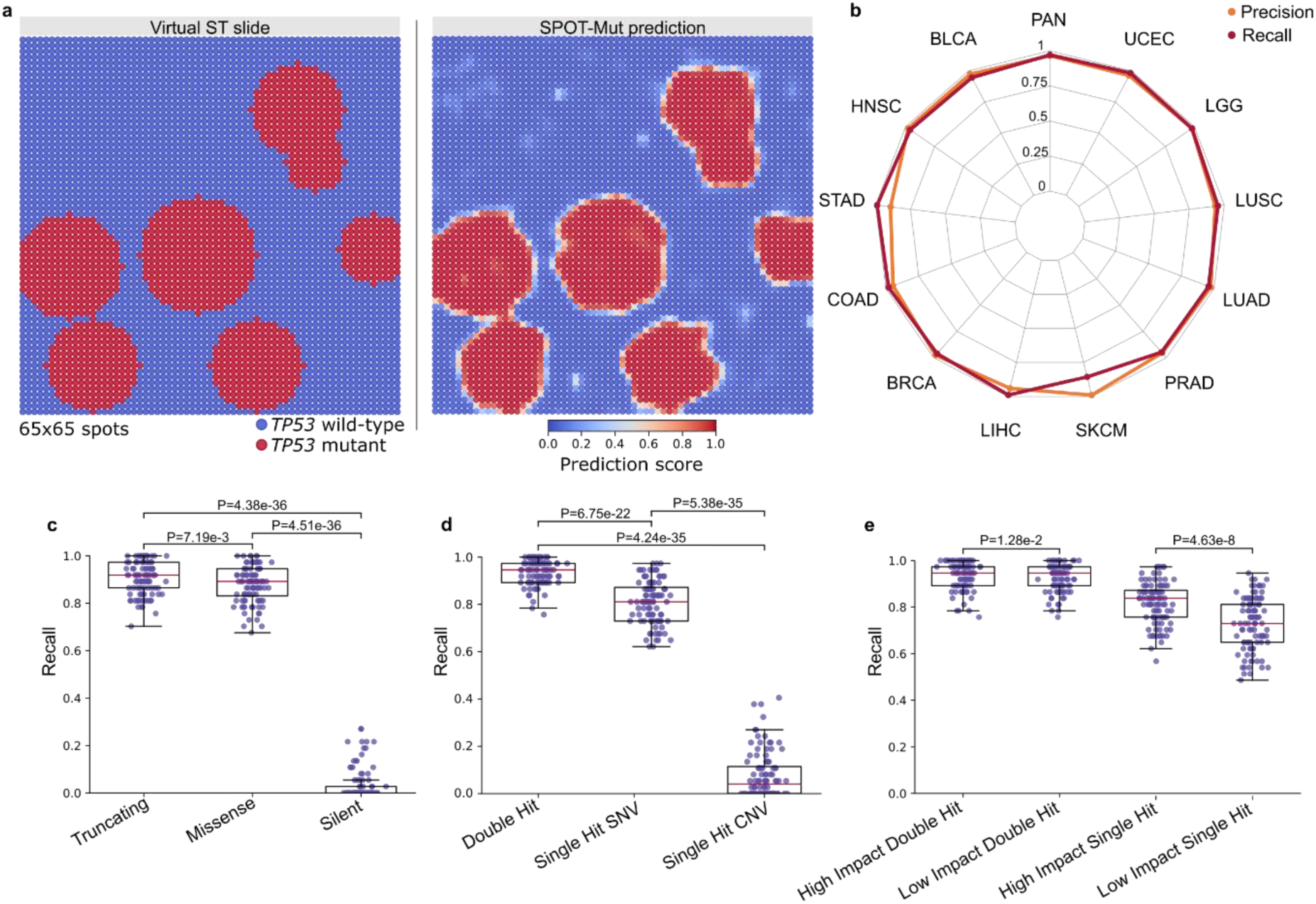
SPOT-Mut *TP53* mutation prediction accuracies in different cancer types and for different mutation types. **a** Example of a simulated ST slide and corresponding SPOT-Mut prediction. **b** Radar plot showing precision and recall values of the TCGA cancer types used to train and test SPOT-Mut, including a pan-cancer category (PAN) that refers to ST slide simulations containing mixed cancer types as spots. Abbreviations are defined in Supplementary Table 1. **c-e** Box plots comparing recall values for 100 clones, simulated from samples with a specific mutation type and/or genomic event. Double hits are defined as samples containing a *TP53* SNV in combination with a copy number loss. High/low impact *TP53* mutations were defined by their PolyPhen-II score (i.e., >0.91 and <0.45, respectively) (33). Box plots represent median values and lower or upper quartiles with whiskers extending to 1.5 times the interquartile range. A one-sided Wilcoxon signed-rank test was used to test pairwise differences between groups.

If these predictions are indeed *TP53* mutation specific, one would expect better prediction accuracies for more damaging mutations (e.g., frameshift or nonsense mutations). To test this hypothesis, we simulated 100 ST slides with a single clone containing only samples with one specific type of *TP53* mutation. Given the relatively low number of samples in each mutation category, we restricted this analysis to the pan-cancer level. Higher recall values were found for truncating mutations (median 0.92) than for missense mutations (median 0.89; *P* = 7.19e-3; one-sided Mann-Whitney *U* test), confirming our hypothesis. As expected, recall values were near zero for silent mutations (median 0; Fig. 2c).

*TP53* is generally known as a haploinsufficient tumour suppressor gene (11), implying a dosage effect with larger expected downstream expression alterations when both alleles are affected (double hits) as compared to single allele alterations (single hits). To determine the sensitivity of the SPOT-Mut model to different allele number alterations, we grouped *TP53* mutated samples in single and dual hits based on their *TP53* copy number state, with *TP53* mutations in copy number neutral samples considered as single hits and in hemizygously deleted samples as double hits. Using the 100 ST slide pan-cancer approach, SPOT-Mut accurately detected samples with a single hit (median recall 0.81). Interestingly, the prediction score was significantly higher in samples with a double hit (median recall 0.95, *P* = 6.75e-22 , one-sided Mann-Whitney *U* test). In contrast, samples that were hemizygously *TP53* deleted in the absence of an SNV were not detected by the model (median recall 0.041; Fig. 2d). In addition, clones with single hit high-impact missense mutations (PolyPhen score >0.91; median recall = 0.84) were significantly better predicted than single hit low-impact clones (PolyPhen score < 0.45; median recall = 0.73; P = 4.63e-8; one-sided Mann-Whitney *U* test) (33). For double-hit samples, the difference between high-and low-impact groups was less pronounced (median recall = 0.95 for both groups; *P* = 1.28e-2; one-sided Mann-Whitney *U* test; Fig. 2e).

### Spatial information is essential for the accuracy of the SPOT-Mut model

To assess the added value of the spatial information to the prediction, we tested two additional, spatial-agnostic models. The first model is similar to the SPOT-Mut model but with an unconnected graph as input, implying the removal of any spatial context. As expected, this unconnected GCNN model showed a worse area under the receiver operating characteristic (AUROC) curve than the SPOT-Mut model (AUROC 0.90 versus 1.00; *P* = 1.6e-174; paired t-test; Fig. 3). Secondly, we trained a generalized linear model (GLM) to predict *TP53* mutation status from RNA-seq. While such a GLM lacks the inherent capability to include spatial data, it has been previously shown to be effective in predicting *TP53* mutations from bulk RNA-seq (10). The GLM was trained and tested on the same TCGA samples as SPOT-Mut, without simulating a spatial relationship between the samples. While this model performed surprisingly better than the unconnected GCNN model (AUROC = 0.94; Fig 3), predictions were still less accurate than the SPOT-Mut model (*P* = 9.61e-202; one-sample t-test). As expected from spatial-agnostic models, these lower accuracies were caused by isolated false positive and false negative spots on the ST slide (Fig 3a). The improved performance over the non-spatial models illustrates the necessity to use spatial relationships to achieve coherent predictions in ST data.

**Fig. 3.**
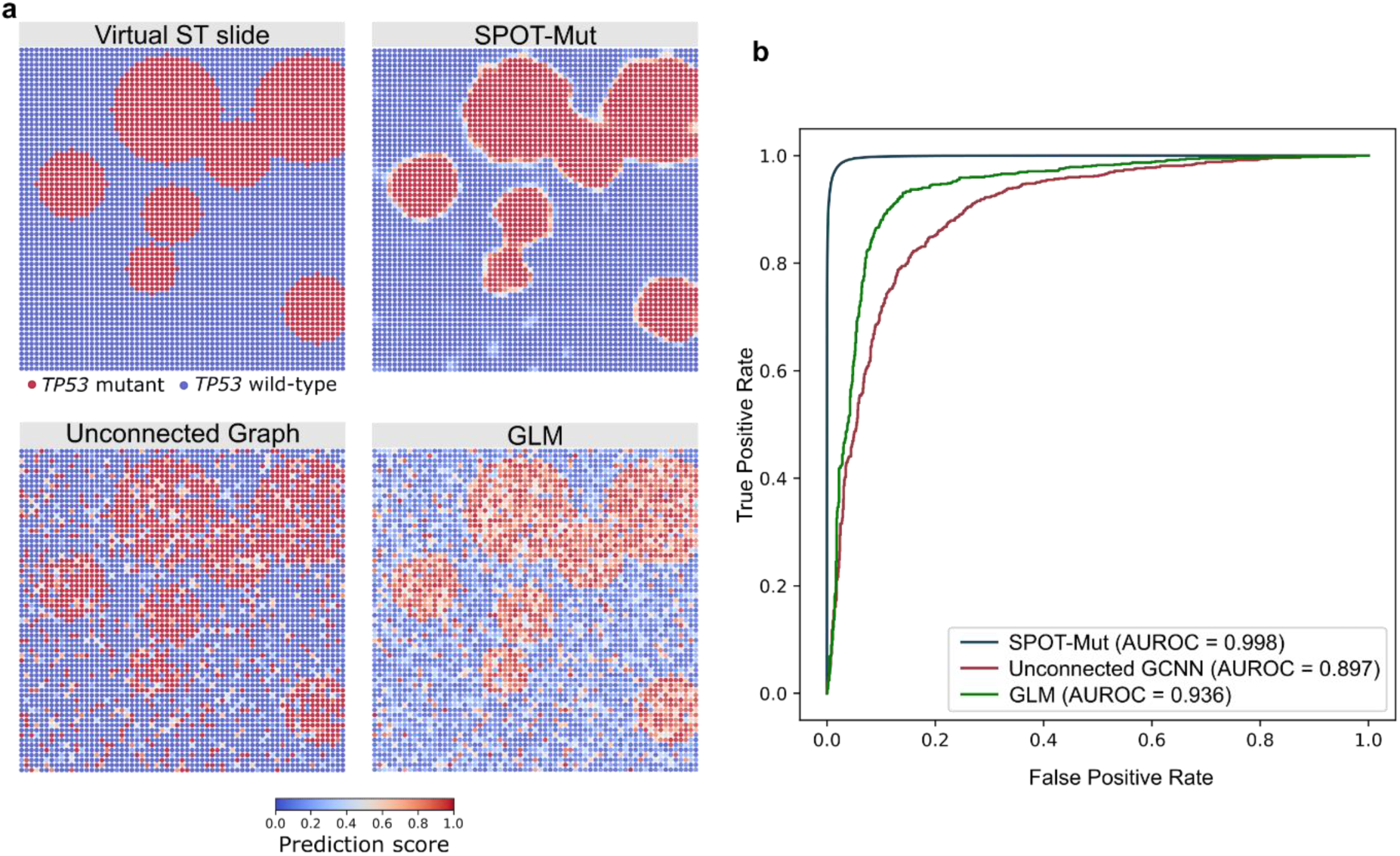
Comparison of *TP53* mutation prediction accuracies between models with and without spatial information 100 virtual ST slides containing *TP53* clones were simulated and predicted using SPOT-Mut and 2 non-spatial models: unconnected GCNN and a generalized linear model (GLM). **a** Representative example of a simulated ST slide and the corresponding mutation probabilities of the three models, as indicated. **b** Receiver Operating Characteristic (ROC) curves comparing the performance of the three different models.

### SPOT-Mut predicts *TP53* mutation status using cancer cell-specific gene expression changes

To test whether the model can be generalized to other data sources, we predicted *TP53*mutation status using cell line RNA-seq data from the Cancer Cell Line Encyclopedia (CCLE) (34). After employing the same virtual ST slide simulation process as described higher, SPOT-Mut identified *TP53* mutated cell lines and recreated the simulated spatial pattern with a performance similar to the weaker performing TCGA cancer types (precision = 0.90, recall = 0. 92; Suppl. Fig. 1a-c).

The generalizability of the model to cancer cells lines implies that SPOT-Mut learns cell-specific gene expression changes, rather than tumour microenvironment (TME) related alterations. The cancer cell specificity was further confirmed by the weaker performance of the model on virtual slides with data from tumours with low purity (< 10^th^ percentile; median recall = 0.70) compared to those with high purity (>90^th^ percentile; median recall = 0.89; *P* = 4.21e-25; one-sided Mann-Whitney *U* test; Suppl. Fig 1d).

### p53-related genes and pathways have the strongest influence on SPOT-Mut predictions

To gain insights in the decision making process of SPOT-Mut, we employed 2 explainable AI (XAI) methods.

Firstly, we applied guided backpropagation (35) to identify which individual features influence the SPOT-Mut predictions. This method calculates attribution values for each feature reflecting its local importance during inference. The *MDM2* gene, encoding the p53 degrader Mmd2, was the feature with the highest absolute attribution values across all spots in all TCGA test datasets (mean attribution = 0.0025). Out of the top 25 genes, 21 were direct p53 targets (36–42) (Fig 4a). When solely focussing on *TP53* mutant spots, *CDKN2A* consistently showed the strongest positive attribution, while *MDM2* exhibited the most negative attribution (i.e., strongest contribution on *TP53* wild-type spots), which are in line with their known opposite effects on p53 signalling (43). A KEGG pathway ranked gene set enrichment analysis (GSEA), using attribution scores as the ranking metric, identified the p53 signalling pathway as the most significantly enriched pathway (NES = 1.88, FDR = 6.7e-8; Fig 4b-c).

**Fig. 4.**
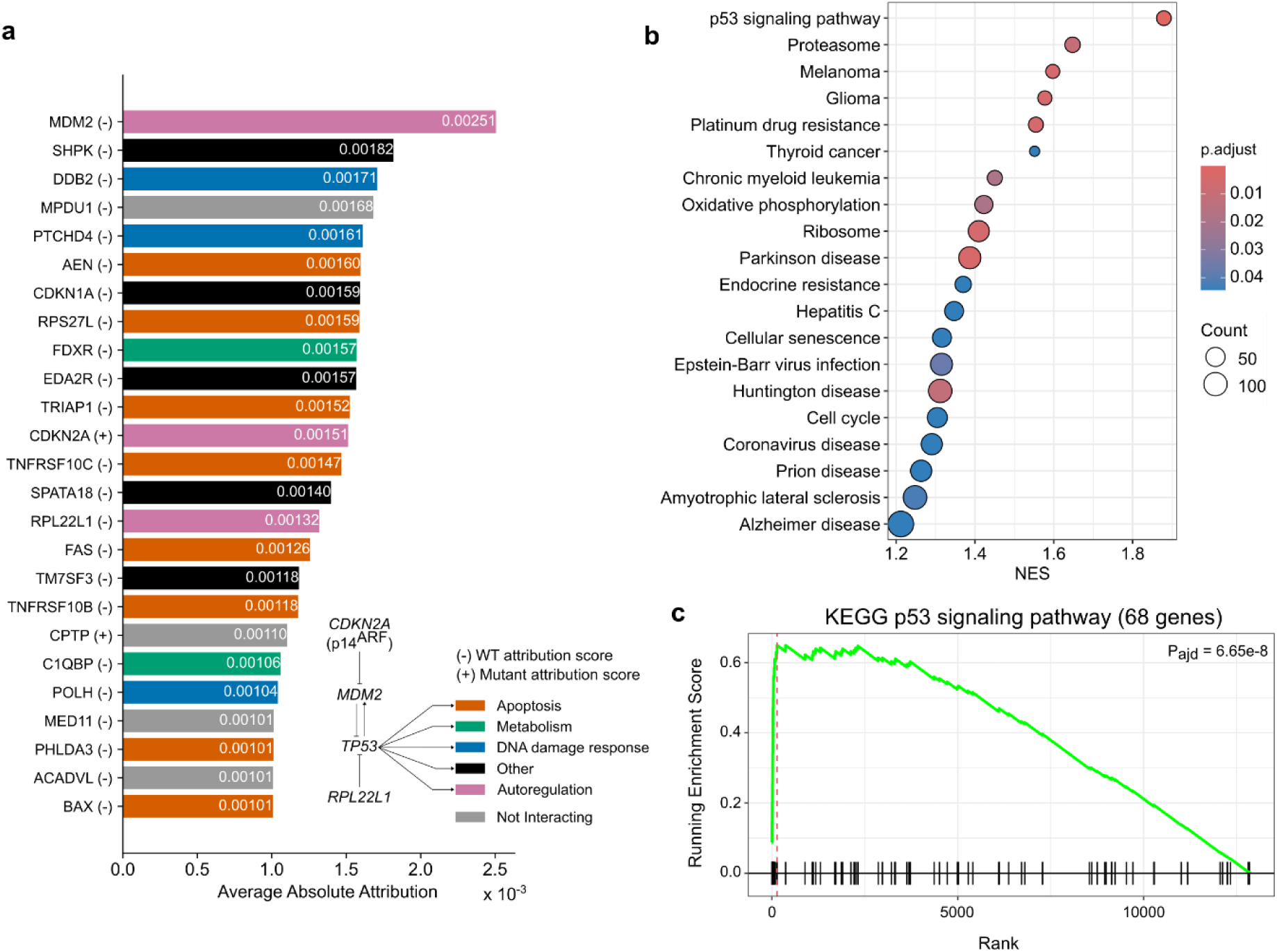
Analysis of genes and pathways explaining the SPOT-Mut predictions **a** Bar plots showing feature attribution scores as calculated using guided-backpropagation. Top 25 genes are shown and ranked based on scores. Known interactions between *TP53* regulators contained in the top 25 are shown on the bottom right. **b** Dot plot showing normalized enrichment scores (NES) and corresponding *p* values for the different pathways as indicated. **c** Running score plot for the KEGG p53 signaling pathway (44).

As a second approach, we aimed to determine which gene sets and/or pathways had the most influence on the prediction accuracy, using Kyoto Encyclopedia of Genes and Genomes (KEGG) pathway gene sets (44). For each pathway, the model performance on the TCGA test dataset was evaluated after removing the involved genes from the datasets (i.e., turning expression values to 0). The resulting AUROC value was then compared to an average baseline AUROC, which was calculated by randomly removing the same number of input genes (100 iterations). Using this approach, we identified genes involved in the *Cell cycl*e (Δ AUROC −1.89e-3) and *p53 Signalling pathway* (Δ AUROC −1.76e-3) and *Pathways in cancer* (Δ AUROC −1.21e-3) as the top 3 contributors to the model performance (Suppl. Fig. 2a).

While both XAI methods clearly prioritized p53-related genes and pathways, the absolute drop in performance of MuT-GCCN was very modest after their removal. This indicates a high level of model robustness, which is further demonstrated by the fact that 12000 random features (i.e., 92% of the 13013 model input features) can be removed without a substantial drop in predictive performance (Suppl. Fig 2b). These results suggest that even for highly sparse data, which is often the case for many ST technologies (45), the model will still perform well

### SPOT-Mut identifies mutant *TP53* subclones in previously published squamous cell carcinoma ST data

Our results demonstrate that SPOT-Mut accurately identifies *TP53* mutated clones in virtual ST data. To test its applicability on real ST data, we downloaded published *Visium V2* data from cutaneous squamous cell carcinoma (cSCC; n = 8; Suppl. Fig. 3a) (2,46) and oral SCC (oSCC; n = 12; Suppl. Fig. 3b) (3), cancer types that are known for their high *TP53* mutation rate (76% for oSCC and up to 95% for cSCC) (47,48). Additionally, we used negative control data obtained from normal skin (n = 12; Suppl. Fig. 4a-g) and atopic dermatitis (AD; n = 7; Suppl. Fig. 4h-s) samples (49). As expected, SPOT-Mut identified *TP53* clones (defined as minimal 2 adjacent spots with a SPOT-Mut score above 0.5) in the majority of oSCC (8/12; 66.7%) and cSCC (6/8; 75%) slides, but not in normal skin epithelium (1/12; 8.3%; Suppl. Fig 4c) or AD (2/7; 28.6%; Suppl. Fig 4o,s) samples. The small clones, with lower probabilities, that were detected in normal skin and AD could indicate the presence of non-tumoural *TP53* clones that have previously been reported in about 5% of aged normal skin epithelium (50,51).

In the cSCC data we observed a striking similarity between the previously identified tumour-specific keratinocytes (TSKs) and the SPOT-Mut prediction score (Spearman ρ = 0.52; *P* < 2.2e-16; Suppl. Fig. 5). These TSKs were characterized by a distinct expression profile, linked to the downregulation of apoptotic and terminal differentiation genes and upregulation of extracellular matrix and cell motility genes. Our results thus suggest a key role of altered *TP53* activity in TSKs.

### Validation of SPOT-Mut predictions in DNA obtained from cSCC sections

The previous analyses demonstrate the applicability of SPOT-Mut on ST data. However, the lack of directly matching DNA sequencing data of the spatially profiled slices limits its validation potential. Therefore we aimed to collect such matched data by serially sectioning 10 µm slices from 3 cSCC samples that were taken *ex vivo;* upon surgical tumour resection in two skin cancer patients. For each tumour, one sample was taken from the tumour core (TC) and for one tumour we also took a sample from the tumour edge (TE). These samples are further referred to as TC1, TC2 and TE2 (number referring to cSCC patient; Fig. 5a).

**Fig. 5.**
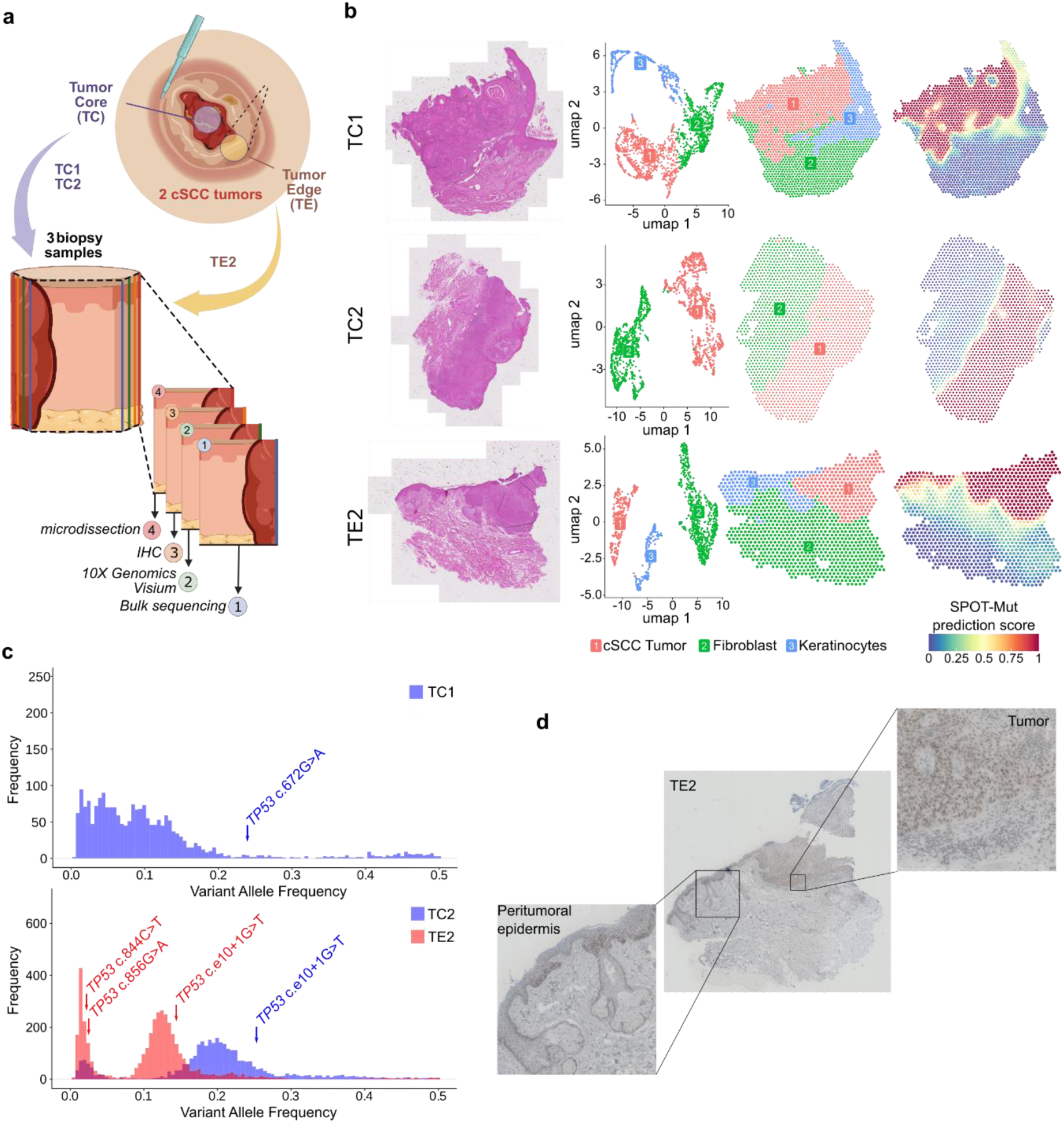
SPOT-Mut predictions and serial slice validations of *TP53* mutations in spatially profiled cSCC samples **a** One tumour edge (TE) and 2 tumour core (TC) samples were obtained from two cSCC tumours (referred to as TC1, TC2 and TE2 as indicated). Consecutive sections were then subjected to whole exome sequencing, ST profiling using the *Visium* platform, p53 IHC or microdissection as indicated. **b** Hematoxylin and eosin staining (H&E), UMAP Leiden clusters (88) and corresponding spatial plots (annotated based on the main cell type) and SPOT-Mut predictions for the four samples as indicated. **c** Variant allele frequency (VAF) distribution of both tumours with TC and TE samples indicated in blue and red, respectively. **d** IHC of TE2 with an anti-p53 primary antibody DO7. Insets in zoom represent epidermis or tumour regions as indicated. Panel a was created in https://BioRender.com

After dermatopathological confirmation of the cSCC diagnosis, one section was spatially profiled using the *Visium V2* ST platform. Tumour cells, keratinocytes and fibroblasts were defined based on their expression profiles (UMAP analysis) and spatial clustering (Fig. 5b). SPOT-Mut demonstrated a clear *TP53* signal in the tumour regions. Interestingly, the epidermis adjacent to the tumour was *TP53*-negative in TC1 but positive in TE2 (Fig. 5b).

DNA from four adjacent sections was isolated and pooled for deep whole-exome sequencing (WES; average sequencing depth 403x). Mutation calling from WES data with GATK Mutect2 (52) identified 64.098 SNVs and short indels across the three samples. Somatic coding SNVs specifically amounted to 2159 for TC1, 3327 for TC2 and 4073 for TE2 (Suppl. Table S2). As expected, the majority of TC2 mutations (2842; 70%) were shared with TE2 and the majority of the mutations had a UV mutational signature SBS7 (53–55), complemented by age-related clock-like signature SBS5 (56) (Suppl. Fig. 6a). In line with the SPOT-Mut prediction, we confirmed the presence of a clonal *TP53* mutation in each tumour (*TP53* c.672G>A in TC1; *TP53* c.e10+1G>T in TC2 and TE2; Fig. 5c). Both mutations were annotated as splice site mutations with high predicted probability of splicing disruption (Δ > 0.90 as calculated using SpliceAI) (57). Interestingly, the variant allele frequency (VAF) of these *TP53* mutations was close to the expected clonal VAF that was calculated based on the SPOT-Mut predicted *TP53* mutant area, further confirming the validity of SPOT-Mut predictions (Suppl. Fig. 6b-e).

Two additional *TP53* missense mutations with low VAF were detected in TE2 but not in TC2 (*TP53* c.856G>A and *TP53* c.844C>T), suggesting a peritumoural epidermal origin, as predicted by SPOT-Mut (Fig. 5b). Similar to the other mutations, a striking similarity was observed between the SPOT-Mut based expected VAF (VAF = 0.025) and the observed VAFs (VAF = 0.025 and 0.022; Fig. 5c; Suppl. Fig. 6b). The results strongly suggest that both mutations originate from the peritumoural epidermis, which was confirmed by additional WES on separately microdissected epidermal, tumoural and dermal regions (Suppl. Fig. 7). Finally, p53 immunohistochemistry indicated elevated nuclear staining for p53 in both the TE2 tumour and the epidermis region, as would be expected from mutant p53 (Fig. 5d). Interestingly, upon further dermatopathological examination, the TE2 epidermis was diagnosed as actinic keratosis, a premalignant skin lesion that has been previously described to often be *TP53* mutated (58–60).

### SPOT-Mut can predict somatic mutations in 40 commonly mutated driver genes

Given the excellent performance of SPOT-Mut in predicting *TP53* mutations, we extended the model to other driver genes. To obtain sufficient and balanced training data, the model was trained on 62 known driver genes (as defined by cancer gene census) that were mutated in minimally 10% of samples and for which sufficient samples (100 per cancer type, minimally 2 cancer types) were available (Suppl. Fig. 8).

Similar to the *TP53* model, high prediction performance was observed on the test data for the majority of genes, with AUROC (median 0.98) and area under the precision-recall curve (AUPRC; median 0.97) being higher than 0.95 for 48 and 40 genes respectively (Fig. 6a-b; Suppl. Table S3). As expected, and in line with our observations for the *TP53* model (Fig. 3), AUROC and other performance metrics values were always higher as compared to non-spatial GLM models (Suppl. Table S3).

**Fig. 6.**
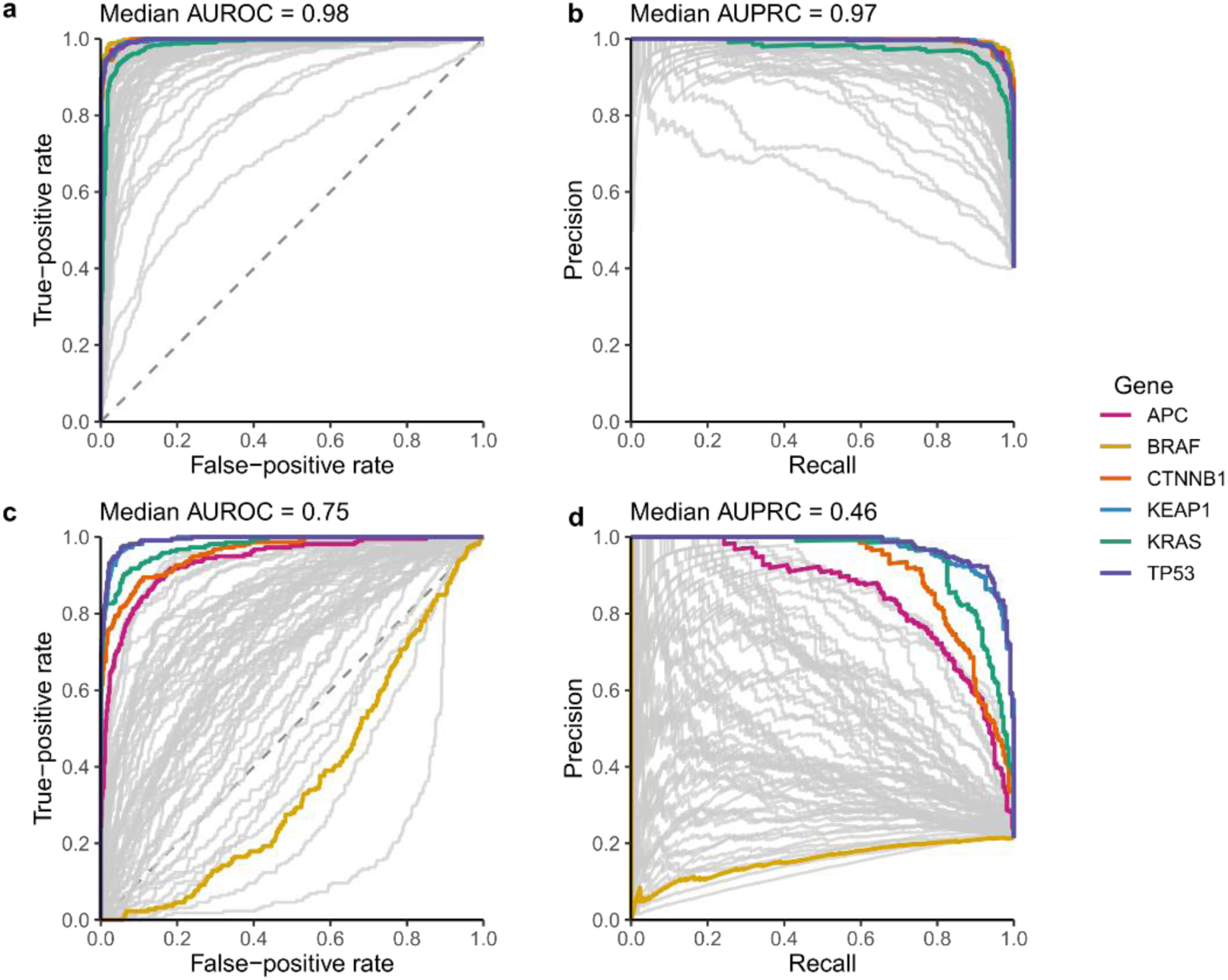
SPOT-Mut prediction performance on 62 common driver genes Receiver operating characteristic (ROC; **a, c**) and precision-recall (PR; **b, d**) curves after training SPOT-Mut models for 62 frequently mutated driver genes and applying the model on simulated ST slides from TCGA (**a, b**) and (c,d) CCLE data. Driver genes that are discussed in main text are highlighted. See Supplementary Table S3 for detailed results.

The generalizability of the 40 well-performing performing models (i.e., with AUROC & AUPRC > 0.95) was further evaluated using simulated ST slides derived from independent pan-cancer cell line data, as described higher (Suppl. Fig. 1). Performance remained high (AUROC and AUPRC > 0.75) for 10 genes, with the models for *TP53* (AUROC = 0.99; AUPRC = 0.98), *KEAP1* (AUROC = 0.99; AUPRC = 0.97), *KRAS* (AUROC = 0.98; AUPRC = 0.94) and *CTNNB1* (AUROC = 0.96; AUPRC = 0.91) performing best (Fig. 6c-d; Suppl. Table S3). Like *TP53*, these results suggest that these well-performing models were highly generalizable and learned cell-intrinsic properties. We could indeed confirm strong enrichments for model-specific pathways when applying the higher described guided backpropagation XAI method (e.g., glutathione metabolism for the *KEAP1* model, NES = 1.86, *P* = 9.49e-8; *MAPK signaling* for the *KRAS* model NES = 1.19, *P* = 1.37e-2; Fatty acid metabolism for the *PTEN* model, NES = 1.65, *P* = 3.34e-5; Wnt Signaling for *CTNNB1* model, NES = 1.41, *P* = 4.13e-5; DNA replication for *RB1*, NES = 2.03, *P* = 6.92e-9; Suppl. Fig. 9).

## Discussion

Spatial transcriptomics is revolutionizing our understanding of tumour biology but inherently lacks information on the upstream somatic driver mutations. We developed SPOT-Mut, a spatially-aware graph convolutional neural network that identifies (sub)clonal mutations in *TP53 ,* the most frequently mutated gene in human cancer (11,13), and extended the approach to 62 other commonly mutated driver genes.

In the absence of large collections of spatial genomics reference datasets, we trained SPOT-Mut on *in silico* generated, virtual ST slides with mutated clones simulated using bulk tumour sequencing data from TCGA. Virtual slides were designed following the default *Visium* layout (65 x 65 spots) and were generated by randomly assigning (bulk) cancer type-specific gene expression data from mutated and wild-type samples to corresponding spots. This methodology implies placing heterogeneous expression data from different patients and tumours in close spatial proximity. Despite its artificial nature, the incorporation of the spatial context during SPOT-Mut training implied that each spot considered the expression information of 24 neighbouring spots (two convolutional layers), which is expected to remove variation that was not directly related to mutation status, the only common spatial context denominator. This likely explains the good generalizability across different cancer types and independent datasets.

The added value of the spatial context to the model’s performance was further demonstrated by the superior prediction accuracy of SPOT-Mut to non-spatial (GLM) models for all evaluated genes. Surprisingly, an unconnected and hence spatially agnostic GCNN model performed worse than the GLM, showing that the GCNN architecture alone does not provide any intrinsic advantage. These results suggest that, while classical linear models are a better alternative for mutation predictions in the absence of any spatial context, GCNNs have the potential to improve performance without relying on post-processing spatial clustering steps, in line with their popularity for deep learning on spatial transcriptomics (61).

A key challenge when developing any novel machine learning approach is the “black box” phenomenon, i.e. does SPOT-Mut indeed predict the expected somatic mutations or is the performance explained and/or biased by hidden, correlated variables? We extensively addressed this challenge for the *TP53* model, by demonstrating *TP53* specificity using three orthogonal approaches: at the genomic, cellular and the molecular layer. Firstly, when evaluating the model on simulated slides that were homogenous for prespecified mutation types, the best recall values were observed for high impact mutations, which are expected to have the largest impact on the *TP53*-specific expression signals. Recall was near-0 for silent mutations, further excluding the possibility that related genomic variables such as tumour mutation burden or mutational signatures confounded our results. Similarly, samples with double hit mutations (*T53* copy number loss in combination with a *TP53* mutation) had higher recall values than samples with a single hit, in line with the expected gene dosage effect of *TP53*, which is known as a haploinsufficient tumour suppressor gene (62). Interestingly, while recall was high (median recall 0.81) for single hit SNV mutations, we demonstrated that the model was largely insensitive to hemizygous deletions in the absence of *TP53* mutations (median recall 0.04). Secondly, when SPOT-Mut was evaluated on virtual slides generated using independent cell line data (34), the accuracy only dropped slightly. Apart from demonstrating the generalizability of our model, this suggests that the model learned *TP53* mutation status from cancer cell-specific expression signals, rather than alterations in the TME, which have been linked to *TP53* mutations previously (63–65). The cell specificity was further confirmed by the better performance in tumours with high purity (high cellularity). Thirdly, two explainable AI approaches pointed to a key role of *TP53* signalling genes and pathways in determining the model’s predictions. Of note, the guided backpropagation approach indicated that Mdm2, which is known as a primary negative regulator of p53 (17–19,66), was the most informative feature for the model. Overall, 21 out of the 25 most informative features were genes directly involved in *TP53* signalling (36–42).

Despite the top contribution of these *TP53* signalling genes and pathways, their absolute contribution was rather modest. This is likely explained by the many existing gene co-expression networks (67,68), with many correlated genes having the potential to provide similar information to the model. This was also suggested by the robustness of SPOT-Mut to sparsity of the input data. Indeed, random removal of genes from the input, even at very large scales, affected the performance only marginally. This robustness to random sparsity is a desirable property as real ST data are sparse by nature. Taken together these results suggest that SPOT-Mut relies on a broad set of biologically meaningful signals rather than a small set of dominant genes for its prediction of *TP53* mutation status.

The *TP53* SPOT-Mut model was validated using an indirect approach on available data and a direct approach on newly generated ST with matched DNA sequencing data. Firstly, when applying SPOT-Mut on previously published non-tumoural (normal skin and AD) and tumoural (cSCC and oSCC) ST datasets, mutation frequencies were in line with expected frequencies from sequencing studies. Interestingly, when applying SPOT-Mut on the cSCC ST samples, the predictions strongly correlated with a cluster of tumour-specific keratinocytes that were identified by the authors of the original study (2). While the data don’t allow us to distinguish between actual subclonal *TP53* mutations versus a strongly altered *TP53* signalling pathway due to other reasons, it does demonstrate how the model can result in novel biological findings. We also validated SPOT-Mut directly using a serial slicing protocol on three samples from two cSCC tumours, with one section being spatially profiled with the *Visium* ST platform and adjacent serial sections being (bulk) DNA sequenced or evaluated using p53 IHC. Clonal *TP53* mutations were predicted in both tumours and confirmed by DNA sequencing. Further, the expected variant allele frequency, as calculated from the SPOT-Mut signal was close to the observed frequency, strongly supporting the identical spatial location of predicted and observed mutations. Unexpectedly, *TP53* mutations were also found in the adjacent epidermal skin to one of the tumours. DNA sequencing confirmed the presence of two missense mutations in this epidermal skin, again with matching observed and expected variant allele frequencies. Interestingly, the adjacent skin was pathologically diagnosed as actinic keratosis, a well-known premalignant skin lesion (69,70).

Training a GCNN on virtual ST slides is in principle possible for any mutated gene of interest as we have demonstrated for 62 driver genes. The main limitation is the availability of sufficient and balanced numbers of wild-type and mutated samples. *TP53* was one of the best performing genes for several reasons. Firstly, its high overall mutation frequency (41% in our dataset) allows for a perfectly balanced training dataset. Secondly, it is a transcription factor, with alterations resulting in direct downstream gene expression changes that are expected to result in optimal model performance. Lastly, it is frequently mutated in many different cancer types, allowing for a good model generalization, which was likely the main reason for the good generalizability of *KRAS* (5 cancer types) and the surprisingly poor generalisability of *BRAF* (2 cancer types, with the majority of mutations derived from thyroid cancer). While generalizability to cell line data could be considered a broad indicator for performance on real ST data, cell line-based simulations were solely performed on a pan-cancer level, posing restrictions on their representability for cancer type-specific models. Evaluating the actual performance on real ST data would require matched ST and DNA sequencing, like we illustrated for the *TP53* model.

SPOT-Mut was developed for the default *Visium* layout but is applicable to data derived from any whole-transcriptome ST platform (e.g., *Visium* HD), as long as the sequencing depth per spot is sufficient. When this is not the case, we suggest merging expression data from neighbouring spots as a preprocessing step before applying SPOT-Mut. While this will reduce the spatial resolution, this is rarely a problem when evaluating multicellular subclonal alterations.

In conclusion SPOT-Mut is a generalizable and validated GCNN that potentiates the reliable spatial prediction of mutated cancer cells in any ST dataset of interest. Adding this hidden genomic information layer has the potential to result in novel biological findings related to the role of *TP53* and other driver gene in carcinogenesis and/or can assist in identifying subclonal tumour regions for targeted microdissection.

## Methods

### TCGA and CCLE data collection

The GDC Data Portal was queried to obtain the download manifest for all open access RNA-seq count files generated from solid tissue samples with the *STAR* aligner by the TCGA consortium (71). With this manifest, the files were downloaded with the GDC Data Transfer Tool (https://github.com/NCI-GDC/gdc-client). From these files, the column containing the unstranded transcripts per million (TPM) counts was extracted for all protein-coding genes. Expression data for mitochondrial genes were excluded due to their order-of-magnitude difference compared to mRNA levels originating from nuclear DNA. Next, for all the downloaded TCGA samples, the mutation annotation format (MAF) file ID’s were collected in a manifest from the GDC Data Portal and downloaded with the GDC Data Transfer Tool. The maf files contain mutation data that can be used to classify samples into the mutated (1) and wild-type (0) categories. Mutations annotated as *silent*, *intron*, *5’ or 3’ UTR* or *3’ or 5’ Flank* were all considered as wild-type.

Cell line mRNA-seq counts were obtained from the DepMap portal release 22Q2 (https://depmap.org/portal; CCLE_expression_full.csv) (72,73). The Log2-transformed TPM expression counts were back-transformed as to obtain raw TMP counts. Additionally, the mutations were obtained from the corresponding mutation file provided on the DepMap portal (CCLE_mutations.csv) and underwent the same filtering steps as the TCGA mutations. Both datasets (TCGA and CCLE) were filtered to include only the genes present in both and then transformed with the natural logarithm to stabilize the variance. Ultimately, RNA-seq data for 1406 CCLE cell lines was used in this study.

13013 genes were selected to be retained in both datasets, and all following datasets used with SPOT-Mut. This is the intersect of genes profiled in TCGA, CCLE and 10X *Visium*, i.e., all relevant datasets for this project. A published cSCC ST sample, deposited in the Gene Expression Omnibus (GEO) under accession number GSM4565825, was used as a representative high-quality 10X Visium experiment (2).

#### Selection of training and test data

For the initial *TP53* model, The TCGA dataset was further filtered down to contain only cancer types with sufficient representable data present. The cut-off was set at a minimum of 250 samples and a *TP53* mutated/wild-type ratio within the interval of [0.1;0.9] to avoid large class imbalances. The remaining dataset consisted of mutation and expression data for 6563 samples from 12 TCGA solid cancer types: BLCA, HNSC, STAD, COAD, BRCA, LIHC, SKCM, PRAD, LUAD, LUSC, LGG and UCEC. This dataset was split into a training and validation set with a 70/30 random split at patient level using the *train_test_split* function implemented in *scikit-learn* v1.9.1 (74). Splitting at patient level made sure samples from the same patient would end up in the same split to not confound the validation dataset with linked data in the training dataset.

The approach was slightly altered for the CGC driver gene screening. TCGA tumour types were retained only if they contained more than 100 samples, of which at least 15 carried a mutation in the target gene, at least 10% were mutation-positive and the final sample collection contained at least 199 samples. For each target gene, the retained samples were divided into a training set (80%) and an independent test set (20%). Splitting was stratified by both tumour type and mutation status to preserve their distributions across the two sets.

Next, the *RobustScaler* scaler was fitted using the TCGA training datasets for each gene (74) and used to scale the training and test datasets.

### *In silico* simulation of spatial transcriptomics

For the initial *TP53* model, a grid of data points was generated, matching the dimensions of a 10X Genomics *Visium* V2 slide, resulting in a 65×65 grid with a total of 4225 spots. In each simulated slide, 7 mutant subclones were modelled by overlaying virtual circles over the grid. The radius of these circles was randomly chosen span between six and eight spots wide for the training dataset and five and twelve for the test dataset. These different size intervals were chosen between training and test to check if the model didn’t learn the size of the circles. Mutated samples were randomly assigned to spots falling within these circles. Normal tissue and wild-type samples were randomly assigned to spots outside the circles. Neighbouring spots are connected to build a grid-shaped undirected, unweighted graph used as input for the graph neural network. To assure each sample is represented on the slide, the random assignment of bulk tissue to grid spots is done in a sequential loop. During each iteration, each sample is assigned only once. If, after all samples are assigned, there are still unoccupied grid spots, the sampling sequence is scrambled before the assignment process resumes. For the CCLE data, grids are simulated using all samples.

For the other driver genes, the approach was slightly different. Because fewer samples were available for some genes, the simulated slides were reduced to a grid of 1024 spots (32 × 32). Each slide contained five mutation-positive clones, positioned randomly and assigned radii between four and seven spots. During training, a new slide was generated from the training data at every epoch, using the epoch number as the random seed. A single fixed slide was generated from the validation data using the same simulation parameters. The same seed was used for this slide for all genes, thereby standardizing the simulated spatial arrangement across models.

### Design and training of the graph convolutional network

A GCNN is a neural network designed to operate on graph-structured data. It can aggregate and transform information from a node’s features, its neighbours and the connections between them to learn representations. SPOT-Mut is tuned to perform node prediction. The GCNN is made up of two subsequent graph convolution layers, implemented in *Pytorch Geometric* (v2.8.0) as *GCNConv* (75). The first *GCNConv* layer receives a matrix of gene expression values for each spatial location or spot, as well as the expression value matrix associated with the surrounding spots. These surrounding locations are defined in the adjacency matrix (A). The values of A were set to zero for the unconnected GCNN. The second *GCNConv* layer performs another convolution sequentially. This output is passed to a multilayer perceptron (MLP) consisting of five subsequent layers, steadily reducing the data dimensionality to a single prediction value which undergoes a sigmoid transformation. The output of each layer in the model is transformed by a rectified linear unit (*ReLu*) activation function. The model contains three dropout layers with a dropout rate of 50%, 20% and 50% respectively placed between the *GCNConv* layers and the first three MLP layers.

The model was trained for 70 epochs using the Adam optimizer. The loss was calculated using *BCEWithLogitsLoss* from PyTorch (version 2.12.0). During training, the loss of the training dataset was monitored. Historically, an epoch is defined as one entire passing of the training data through the algorithm. In this context, the classical definition of an *epoch* should be adapted, as most samples are reused multiple times to create one simulated ST slide. Thus, we define one *epoch* as one entire passing of all simulated ST slides through the GCNN. The order of slides passing through the model is random. This means the model encounters each TCGA sample multiple times in each epoch. Each time, the sample will have different neighbours, which changes the context. Eventually, the model obtained in the final epoch had the lowest average loss score on the validation slides and was chosen as the final trained model to be used for further analyses. A binarization threshold was identified by computing the mean misclassification error (MSE) across cutoff values in increments of 0.01 on all validation data, and selecting the cutoff at which the MSE was minimized. Every prediction value above this cutoff ought to be considered as a mutated spot, while wild-type spots would result in a score lower than the cutoff.

### SPOT-Mut performance evaluation

Since the assignment of mutated clones to the simulated ST slide is random, 100 different slides with random clone layouts were simulated to evaluate model performance. Across these slides, precision and recall values were calculated. This was done separately using (I) validation samples from each included TCGA cancer type, (II) all TCGA validation samples (pan-cancer) and (III) the CCLE data. To evaluate the impact of spatial connections on model performance, unconnected ST slides were generated from the same 100 pan-cancer ST simulations, with edge connections set as an empty list.

To evaluate the impact of specific mutation types on model prediction, 65×65 ST simulations were generated with the a single central clone with 7 spot diameter. This clone was occupied by spots containing TCGA validation samples with a specific mutation type. Recall values for only the clonal spots were calculated. This was repeated 100 times for each mutation type.

### TCGA purity estimates

Purity estimates of TCGA samples were obtained using the mean purity estimations from 5 different methods (76). Missing (NaN) values were disregarded. Samples above the 90^th^ percentile were classified as *high-purity*, samples below the 10^th^ percentile were classified as *low-purity*.

### GLM

An binomial elastic net penalized logistic regression model was fitted to the TCGA data to compare performance against the GCNN. The same data used to train and validate SPOT-Mut was used to train and validate this GLM. The *RobustScaler* scaling was not applied and no ST slides were simulated as the GLM doesn’t use spatial information. First, the optimal lambda value was determined through 10-fold cross-validation using the *cv.glmnet* function (glmnet 4.1-8 R package) with an alpha value of 0.2. Next, the lambda value from the fold which produced the lowest mean cross-validated error (*lambda.min*) was provided to fit the final GLM to with the training data using the *glmnet* R function from the same glmnet R package (77–79). On the validation dataset, the ROC curve and AUROC value was determined to evaluate the performance. One-sample Mann-Whitney U test was performed to check if the AUROC value differed from the distribution of AUROC values obtained with SPOT-Mut pan-cancer validation virtual slides.

### Public Spatial Transcriptomics datasets

For all real ST data used, raw count tables and supplement information, such as images, were downloaded from the databases listed in their accompanying publication. These datasets were loaded into an R Seurat (v5.1.0) object and normalised with *NormalizeData* with *LogNormalize* normalization method and *scale.factor* set to 1e6. *ALRA* (v0.0.0.9000) (80) was used to lower the data sparsity by imputing missing count values. Next, the ALRA count matrix was batch corrected using *ComBat* (81), from the *sva* R package (v3.50.0) with the full TCGA dataset as reference batch. This means the TCGA dataset remains unaltered while the ST count matrix is collectively batch corrected. Finally, the batch corrected count matrix was scaled and centered using *RobustScaler* function, implemented in the *scikit-learn* (v1.9.1) python package and fitted on the TCGA training dataset. After defining the connections between neighboring spots to generate the graph structure, the data is ready as input to SPOT-Mut. Our in-house generated ST experiments were processed the same way starting from the 10X Genomics *SpaceRanger* output.

SPOT-Mut prediction values for each cSCC ST slide were correlated using the spearman correlation method to the gene expression programs from a gene set characterizing tumour-specific keratinocytes (TSK). The gene expression program values were calculated with Seurat (v5.5.0) *AddModuleScore* with default parameters (82,83). TSK Correlations were only computed for cSCC ST samples because this expression signature is specific for cSCC (84).

### Guided backpropagation

Feature attribution scores for individual genes were calculated using the guided backpropagation method (35) from the *Captum* library (85) as implemented in the *explain* module of *Pytorch geometric* (v2.8.0) (86,87). By the design of the graph convolutional neural network, nodes more than two connections away don’t influence the node on which the model is generating a prediction. Thus, only the local, reachable, neighbourhood was considered for each node, after confirming that the inclusion of farther nodes don’t influence the attribution scores five different central nodes. For each node in the test dataset, guided backpropagation was applied to obtain feature attribution scores. These were averaged across all nodes of a simulated ST slide. Across all positive and negative nodes separately to obtain attribution scores specific to both prediction categories (i.e., mutated vs wild-type). Absolute values were taken to rank the features according to importance for the values that were averaged across all nodes.

### Gene set enrichtment analysis

The curated human Kyoto Encyclopedia of Genes and Genomes (KEGG) pathway gene sets were obtained (44) against which ranked gene set enrichtment analysis (GSEA) was performed using the *gseKEGG* function from *clusterProfiler* v4.14.0 in R. *pAdjustMethod* was set to ‘fdr’.

### Feature ablation analyses

The curated human Kyoto Encyclopedia of Genes and Genomes (KEGG) pathway gene sets were obtained (44). For each KEGG pathway gene set, the corresponding inputs for SPOT-Mut were set to zero for each cancer type in the test dataset and the AUROC of the prediction result was recorded. To compare, a random set of genes with equal size as the KEGG pathway gene set was also removed from the input (i.e., set to zero). This process was bootstrapped 100x and the average AUROC value was taken. The impact of KEGG pathway gene set removal was then defined as the AUROC difference between random removal and KEGG pathway gene set removal, averaged across all TCGA cancer types in the test dataset.

To estimate the network robustness, random genes were dropped from the input of the TCGA test dataset by setting the expression value of the gene to 0 for each spot on the simulated slide. The number of genes varied from 0 to 13000 in steps of 1000. This operation was repeated 100 times, and the average AUROC value was calculated across those repetitions. This was compared for both an input graph with and without connections between the nodes. Additionally, genes were stepwise (0 to 13000 in steps of 1000) removed from the input according to their rank based on attribution score. The AUROC value after removal was noted and compared to random removal of genes.

### Spatial transcriptomics

To validate SPOT-Mut predictions, two cSCC tumour core and two tumour edge samples were resected from two different patients at the Ghent University Hospital, Belgium. The collection and use of human study samples were approved by the Ethics Committee of Ghent University Hospital (approval number B6702023000083). All patients provided written informed consent. Samples from the tumour core and tumour edge region were taken using 6mm biopsy punches, fixed in 10% NBF and subsequently processed into FFPE blocks.

These FFPE sample blocks underwent a parallel sectioning protocol using the HistoCore Multicut microtome (Leica Biosystems). First, sections were taken with a thickness of 10µm and consecutively used for RNA QC (4 sections) and DNA sequencing (see next). The next suitable section with a thickness of 5µm was used for spatial transcriptomics with 10X Genomics FFPE *Visium* v2.0 platform, using the 10X Genomics *Visium* CytAssist FFPE workflow on a Spatial Gene Expression Slide v2 (6.5mm). Libraries were sequenced on an Illumina NovaSeq 6000 (S4 flow cell, v1.5 reagent kit, 200 cycles) in paired-end, dual-index mode using the following sequencing configuration: Read 1, 28 cycles; i7, 10 cycles; i5, 10 cycles; Read 2, 90 cycles, with a 1% PhiX control spike-in. Sequencing was performed at the VIB Nucleomics Core. 10X Genomics *SpaceRanger* 3.1.2 was used to process the spatial sequencing data and align the reads to GRCh38. A small number of spots located outside the tissue boundaries were retained after *SpaceRanger* processing and were removed following visual inspection in the 10X Genomics *Loupe Browser*. Finally, the data was loaded as a Seurat object in R to perform basic QC by checking for outliers in the number of features and reads per spot. Spots with less than 100 different features were excluded. This amounted to the removal of 8 spots across all 4 ST slides. Due to excessive tissue shearing and lack of visible tumour, the tumour edge sample of patient 1 was disregarded from further analysis. Data preprocessing for SPOT-Mut prediction was performed as discussed earlier.

### UMAP Clustering

UMAP-based clusters were determined using *Leiden* algorithm implemented in *FindClusters* function from Seurat (v5.5.0) in R (83,88). The *resolution* parameter was optimized to resolve distinct cell populations corresponding to histologically defined skin compartments, including tumour cells, fibroblasts and keratinocytes. For TC1 and TE2, the resolution was set at 0.15, whereas for samples TE1 and TC1, a resolution value of 0.1 was applied. tumour

### Immunohistochemistry

FFPE tissue sections from TC1 and TE2 were deparaffinized and rehydrated. Endogenous peroxidase activity was blocked, followed by heat-induced antigen retrieval in citrate buffer (pH 6.0) for 10 min. Sections were incubated with anti-p53 primary antibody (DO-7, 1:100; Cell Signaling Technology, Cat# 48818S) and an HRP-conjugated secondary antibody. Immunoreactivity was visualized using a DAB Substrate Kit (Abcam, Cat# ab64264). Slides were counterstained with hematoxylin, dehydrated, and mounted for microscopic examination.

### DNA sequencing and read alignment

The cSCC tumour samples underwent paired-end whole-exome sequencing (150bp read length) at the NXTGNT, a Ghent University core facility. For this experiment, four subsequent sections were obtained prior to the ST section. These were pooled and DNA was extracted using the QIAamp® DNA FFPE Tissue Kit and Deparaffinization Solution. Sequencing was performed on the AVITI24 sequencer by Element Biosciences following the Trinity library preparation protocol for whole-exome sequencing using the Twist Whole-exome 2.0 panel from Twist Biosciences. The resulting fastq files were processed following the GATK best practices workflow for short somatic variant discovery (52,89,90). First, the sequencing data was aligned to the UCSC hg38 human reference genome (91) using BWA-mem2 (v2.3) (92) for each sequencing run separately. Read groups were assigned following the GATK recommendations (93). Next, the aligned BAM files for each run were merged per sample using *SAMtools* (v1.21) (94). PCR duplicates were marked with *GATK* (v4.6.2.0) *MarkDuplicates*. Base Quality Score Recalibration (BQSR) is a preprocessing step to detect systematic errors in base quality scores. This step was performed with the GATK tools *BaseRecalibrator* and *ApplyBQSR*. The *af-only-gnomad.hg38.vcf.gz* file, provided by the GATK Resource Bundle (95), was used in the --known-site argument of BaseRecalibrator.

### Somatic variant calling

GATK (v4.6.2.0) *Mutect2* was used for short variant discovery in tumour-only mode, as no normal samples were available. The --callable-depth argument was set to 20, the same Gnomad (96) *af-only-gnomad.hg38.vcf.gz* file was used for the --germline-resource option and *1000g_pon.hg38.vcf.gz*, also provided in the GATK Recourse Bundle, was used for the -- panel-of-normals option. Resulting variants were filtered using GATK *FilterMutectCalls* following the Best Practices recommendations. To further remove unreliable variants, only variants in canonical chromosomes with a coverage of at least 10x were kept. Additionally, variants with a population frequency higher than 1% according to dbSNP TOPMED were removed (97). Variants were annotated using GATK Funcotator, using the Data Source v1.7.20200521s as reference database. The resulting mutations are reported in Supplementary Table S2. Coding SNVs were determined as single-nucleotide mutations classified as Missense_Mutation, Nonsense_Mutation, Splice_Site, Nonstop_Mutation, Silent START_CODON_SNP, DE_NOVO_START_IN_FRAME, DE_NOVO_START_OUT_FRAME.

### Functional annotation

Finally, functional annotations were performed with GATK *Funcotator*. All *TP53* mutations were uploaded to the *SpliceAI* website (Dec 14 2025 version) for functional evaluation (57). All settings were kept default. *SpliceAI* Delta scores were noted.

### Mutational signatures

The trinucleotide profile for each tumour was matched against known mutational signatures from COSMIC v3.3 using the R implementation of *SigProfilerAssignmentR* v1.1.3 for exome sequencing data. *Artifact_signatures*, *Lymphoid_signatures*, *AA_signatures* and *Colibactin_signatures* signature groups were excluded. Otherwise, default parameters were used.

### Expected allele frequency calculations

For samples with *TP53* mutations identified in the WES data (TC1, TC2 and TE2) the expected allele frequency (AF) was calculated for the SPOT-Mut-predicted *TP53* mutant regions. First an image of the prediction was opened in Inkscape. A shape was manually drawn around the predicted *TP53* mutant region. The surface area of this shape was calculated using the Inkscape path measurement function. The same operation was performed to delineate the surface area of the full tissue slide. The expected AF was calculated as:

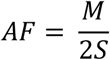

With *M* the surface area of the *TP53* mutant region and *S* the total tissue area. Both units are pixels squared (px²).

### Manual microdissection

Parallel FFPE tissue sections from TC1, TC2, and TE2 were mounted on non-adhesive microscope slides (Thermo Scientific, Cat# 630-1985) using the Epredia™ HM 355S Automatic Microtome. A total of 11, 15, and 11 consecutive sections with a thickness of 10µm were collected for TC1, TC2, and TE2, respectively. Subsequently, the regions of interest were outlined using a permanent marker. TC1 was divided into three regions (tumour, epithelium (epi), and normal dermis), TC2 was divided into three regions (tumour 1, tumour 2, and normal dermis), TE2 was divided into three regions (tumour, epithelium (epi), and normal dermis). The regions were separated using a scalpel, scraped off the slides, and collected in separate 2 mL tubes. DNA extraction of the samples was performed according to the QIAamp DNA FFPE Tissue Kit protocol (QIAGEN, Cat# 56404). WES library preparation was performed with the Agilent SureSelect Human All Exon v6 kit and sequencing was run on the Illumina Novaseq X plus platform with a target coverage of 200x. Due to low input concentrations and high DNA degradation, the effective coverage was lower. Reads were aligned to the human hg38 reference genome using BWA-mem2 (v2.3) (92). Duplicate reads were called with GATK (v4.6.2.0) MarkDuplicates. Coverage at *TP53* mutant sites of interest was evaluated by loading the aligned BAM files into IGV (v2.19.6) and observing the number of reads carrying the mutant and wild-type alleles.

## Data and code availability

Code used to produce the results reported in this manuscript are available from https://github.com/CCGGlab/SPOT-Mut. SPOT-Mut models for all 62 evaluated genes are available to explore predicted somatic mutations in newly generated ST data via an interactive web application that is accessible at https://ccgg.ugent.be/shiny/spot-mut/. GCNN model weights and data scalers can be downloaded from this application.

## Acknowledgements

This work was supported by the Kom op tegen Kanker (Stand up to Cancer), the Flemish cancer society (TL; STI.VLK.2022.0002.01), Research Foundation – Flanders FWO (TL; FWO.3F0.2022.0045.01 and JVdE; FWO.OPR.2025.0043.01) and Cancer Research Institute Ghent (IH, JVdE; partnership grant).

## Author contributions

J.V.d.E and T.L. designed the study and drafted the manuscript. T.L. programmed the model, performed the (bio)informatics analyses. I.H. was responsible for medical diagnosis, inclusion and supervised sampling of the skin cancer patients. S.H. was responsible for DNA extraction, concentration measurements and integrity checks. S.D.H. and E.P. performed pathological inspection of the histology images. S.W. performed the p53 IHC. E.D.M and F.V.N. performed the whole-exome sequencing. E.V.H was responsible for the spatial transcriptomics experiments.

## Additional information

The author(s) declare no competing interests.

