## Supplementary Figures for "Predicting clonal somatic mutations from tumour spatial transcriptomics data using a graph convolutional neural network"

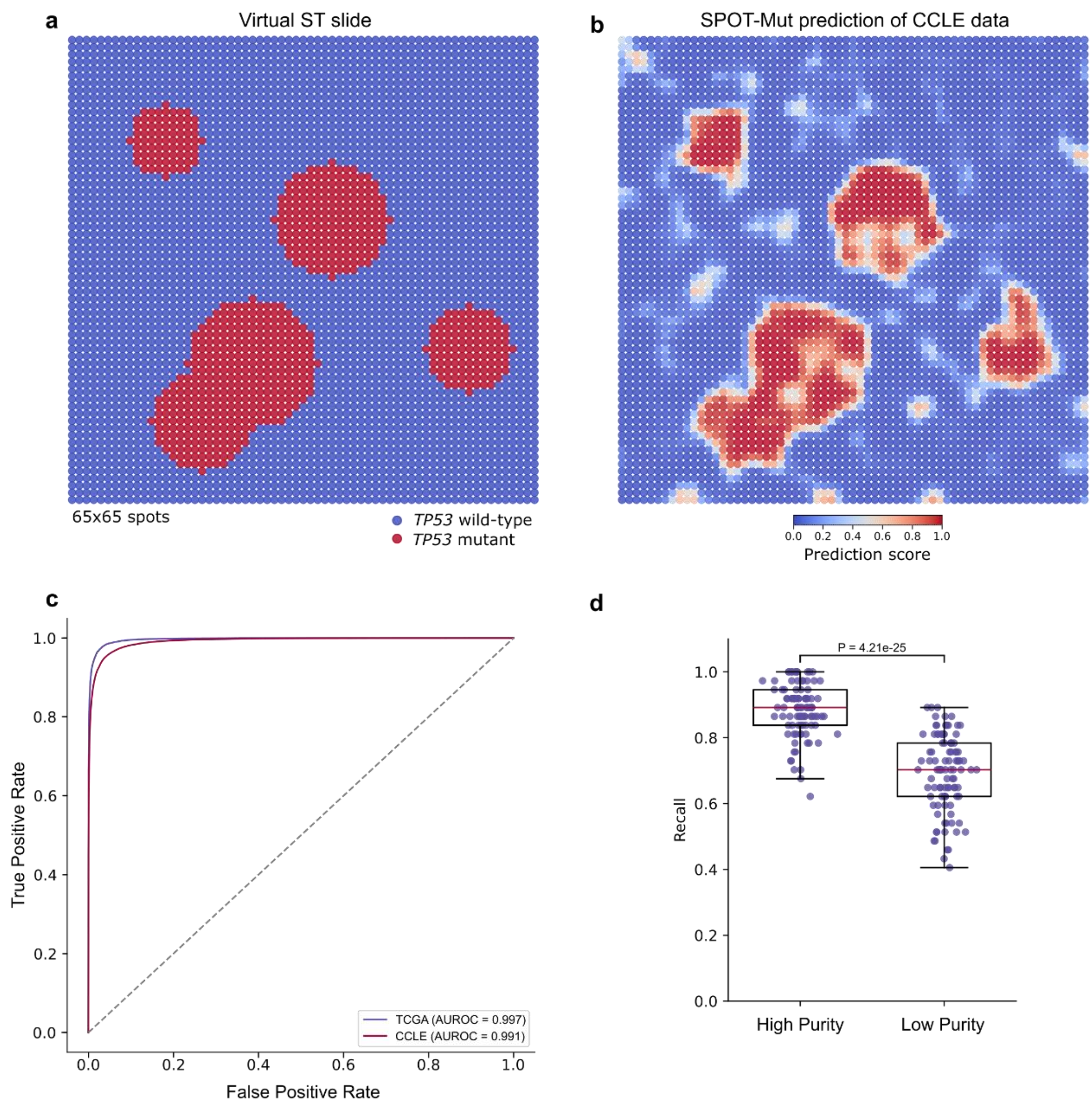


**Supplementary Fig. 1 SPOT-Mut predictions on virtual ST slides simulated from CCLE data**

**a-b** Example of a virtual ST slide simulated with cell-line expression data obtained from the Cancer Cell Line Enyclopedia (CCLE) database (72), and corresponding SPOT-Mut prediction. **c** Receiver Operating Characteristic (ROC) curves comparing the performance of the SPOT-Mut on 100 simulated CCLE slide versus the TCGA pan-cancer (PAN) data. **d** Box plots comparing recall values for 100 simulated clones populated with either high- ($>90^{th}$ percentile) or low ($<10^{th}$ percentile) purity TCGA tumor samples. Box plots represent median values and lower or upper quartiles with whiskers extending to 1.5 times the interquartile range. A one-sided Wilcoxon signed-rank test was used to test pairwise differences between groups.

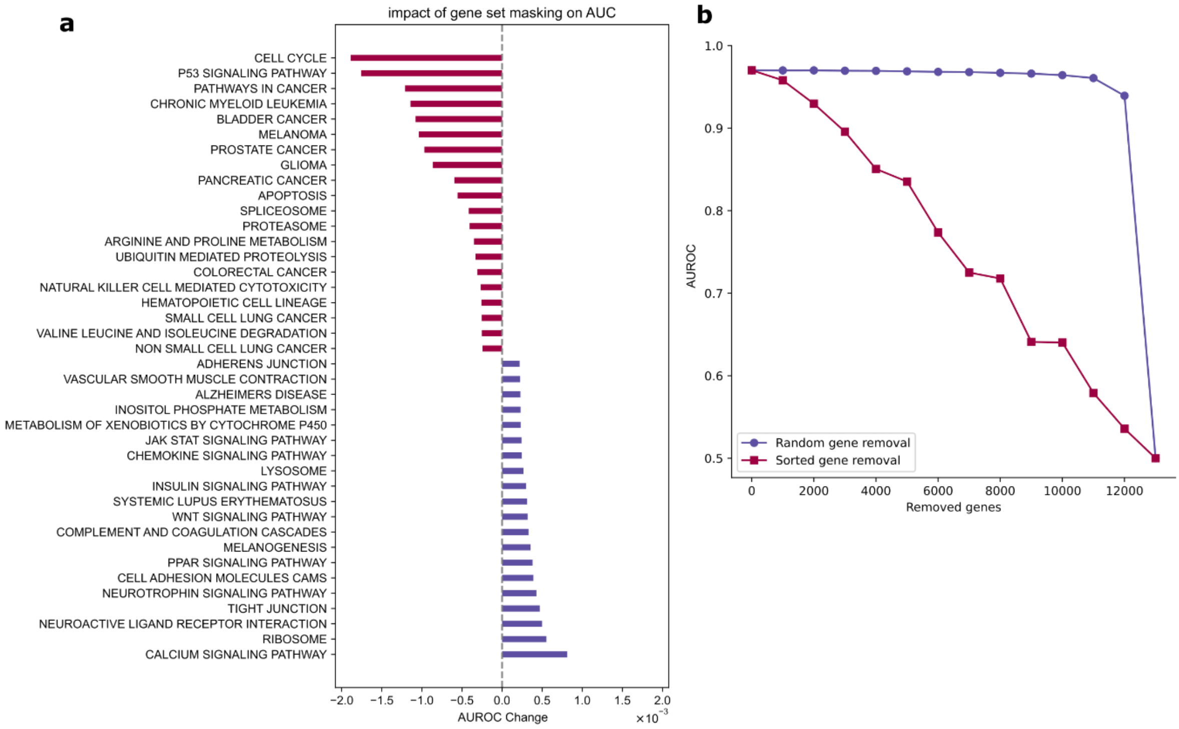


**Supplementary Fig. 2 Feature attribution assigned to KEGG p53 signalling pathway and gene ablation effects on AUROC**

**a** Bar plot showing changes in AUROC after input feature removal of KEGG gene sets.The zero line was calculated as the average AUROC after 100 bootstrap iterations of random input feature removal of the same number of genes as the gene set. Top and bottom 25 pathways shown and ranked according to AUROC change. **b** Comparison of AUROC values between random gene removal with versus in order of guided backpropagation feature attribution score from the SPOT-Mut input.

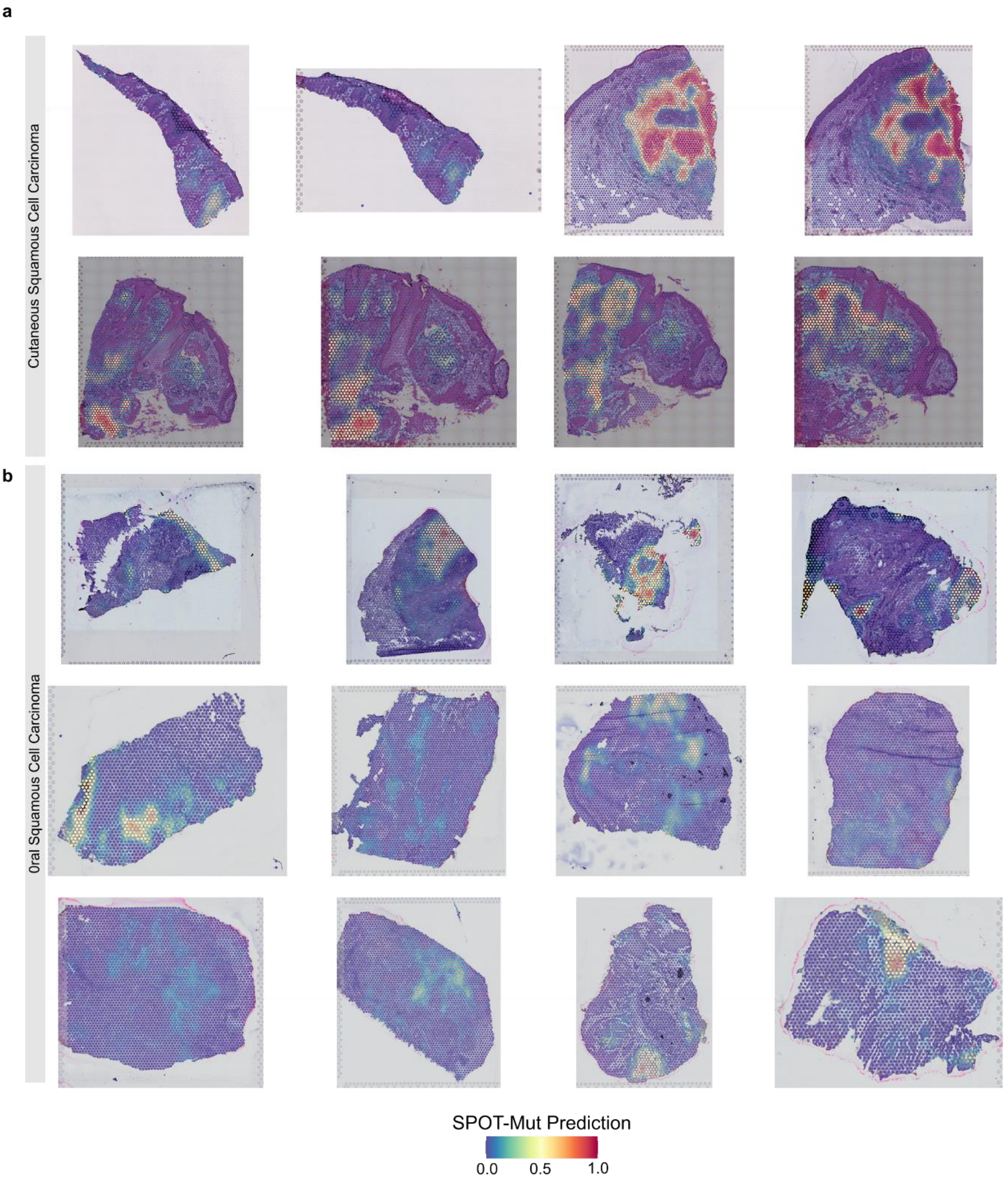


**Supplementary Fig. 3 SPOT-Mut predictions of cSCC and oSCC spatial transcriptomics datasets**

SPOT-Mut predictions, ranging from 0 to 1, visualized on the histology of **a** 8 cSCC spatial transcriptomics (ST) samples (2) and **b** 12 oSCC ST samples (3). A *TP53* clone is detected when ≥2 spots within a single slide have a SPOT-Mut prediction score above 0.5.

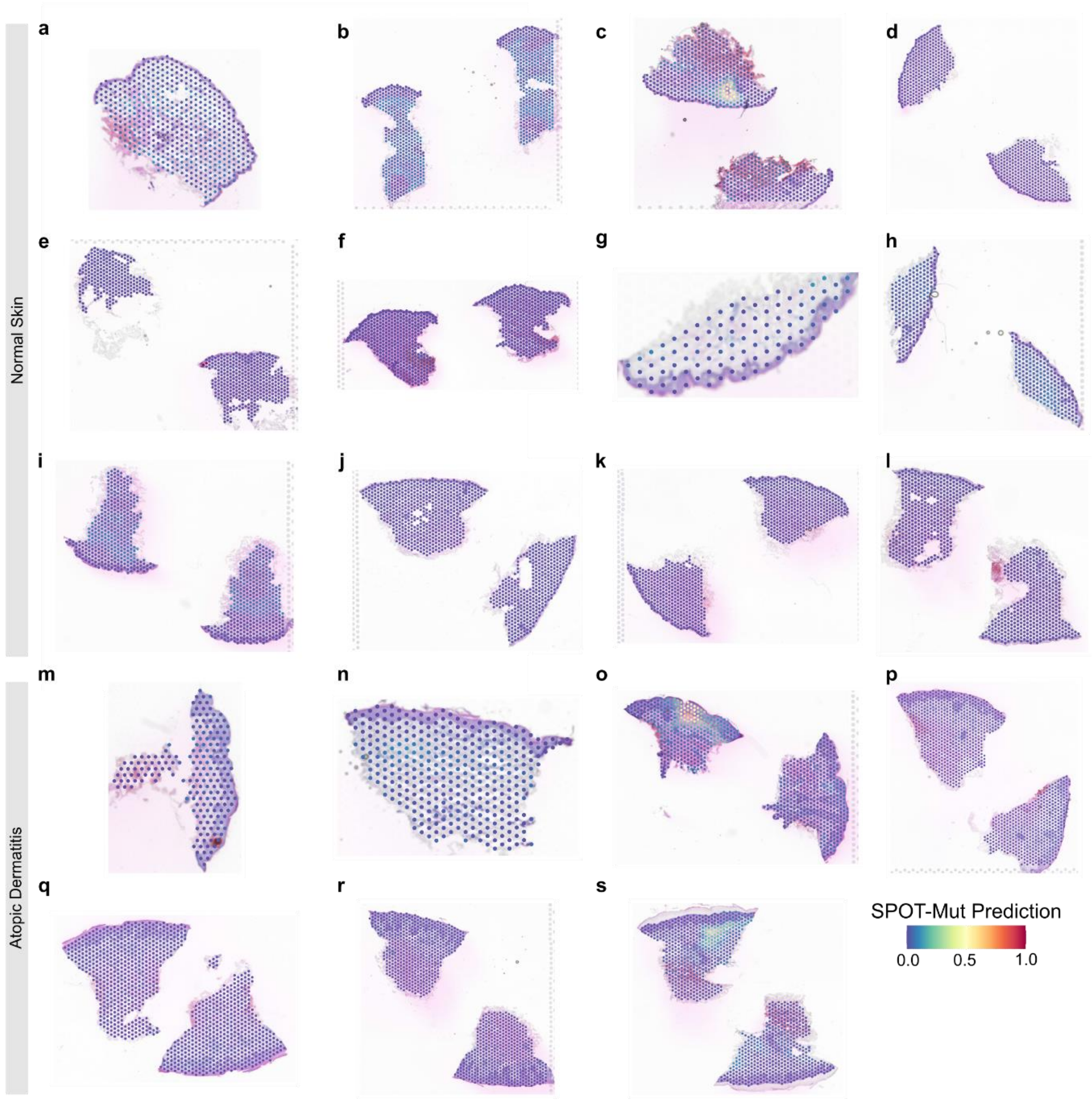


**Supplementary Fig. 4 SPOT-Mut predictions of normal skin and atopic dermatitis spatial transcriptomics data**

SPOT-Mut predictions, ranging from 0 to 1, visualized on the histology across **a-l** 12 spatial transcriptomics (ST) samples derived from normal skin. **m-s** 7 ST samples derived from atopic dermatitis (AD) skin (49). A *TP53* clone is detected when ≥2 spots within a single slide have a SPOT-Mut prediction score above 0.5.

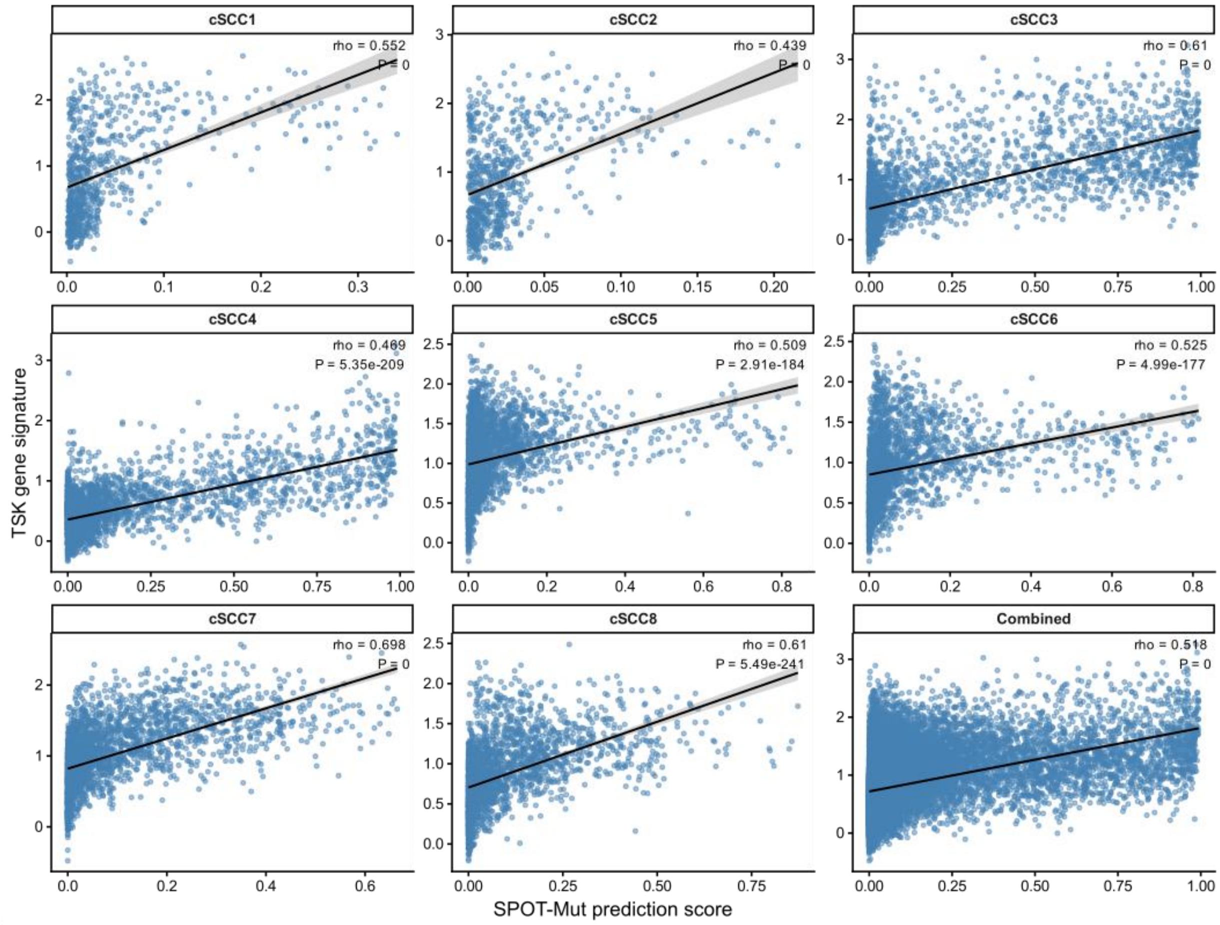


**Supplementary Fig. 5 SPOT-Mut prediction score correlation with TSK expression module in cSCC**

Spearman rank correlations between SPOT-Mut prediction score and tumor-specific keratinocytes (TSK) expression module score as determined by the Seurat *AddModuleScore* function. The correlation is shown for 8 cSCC samples separately and all ST samples combined in the final panel (2).

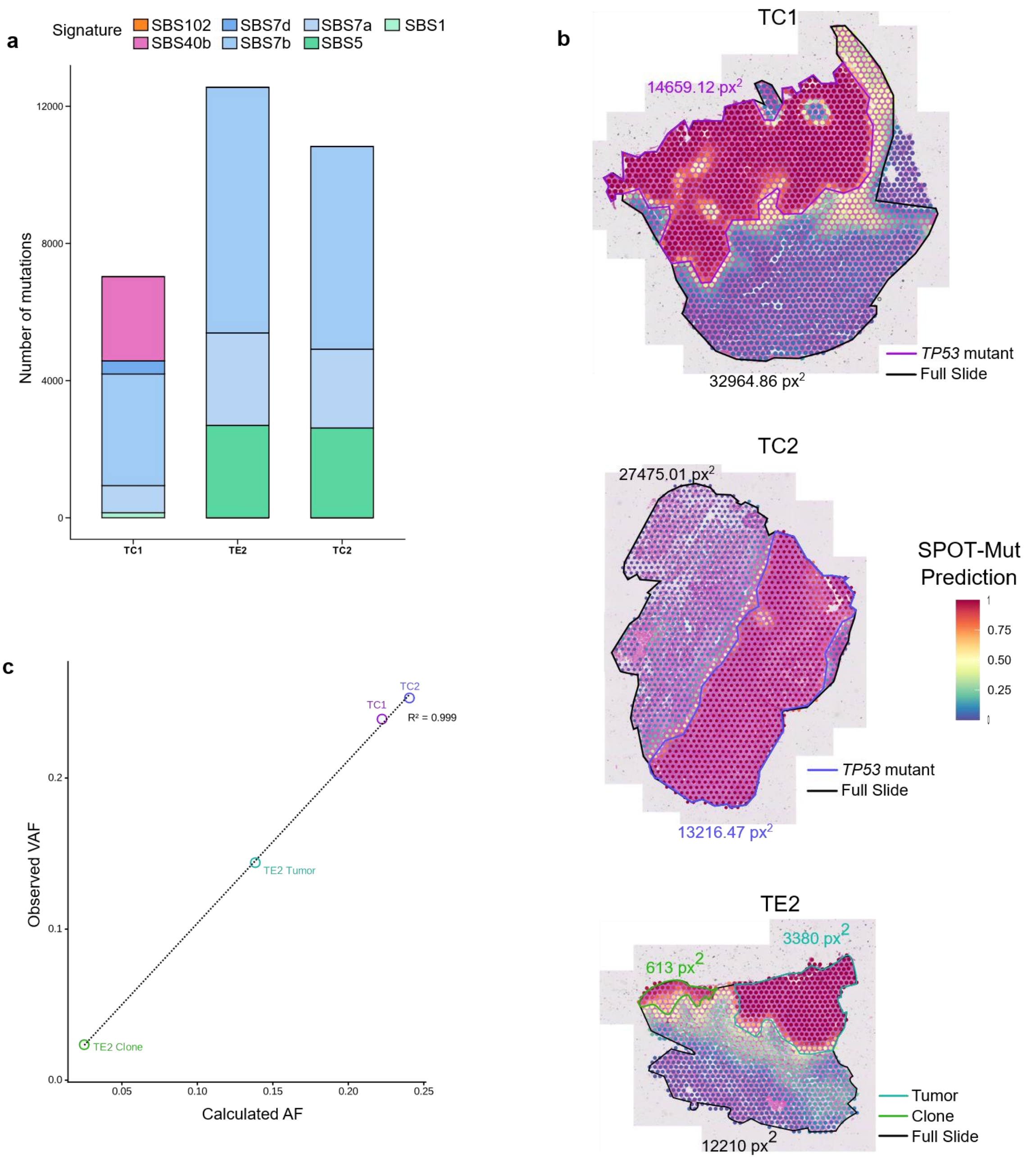


**Supplementary Fig. 6 Mutational signatures and delineation of SPOT-Mut predicted *TP53* mutant areas**

**a** Absolute mutational signature contributions based on mutations identified from whole-exome sequencing (WES). SBS signatures are derived from the COSMIC v3.3 definitions (98). **b** Correlation between observed variant allele frequency (VAF) of *TP53* mutations and calculated allele frequency (AF) estimated from the SPOT-Mut-predicted *TP53*-mutant area within the corresponding sample. Colors correspond to delineated areas shown panels c-d. **b** Delineation of the SPOT-Mut predicted TP53 mutant regions. The area of each delineation is indicated as px². Expected VAF was calculated as half of the fraction of a *TP53* mutated area relative to the total tissue area ("Full Slide").

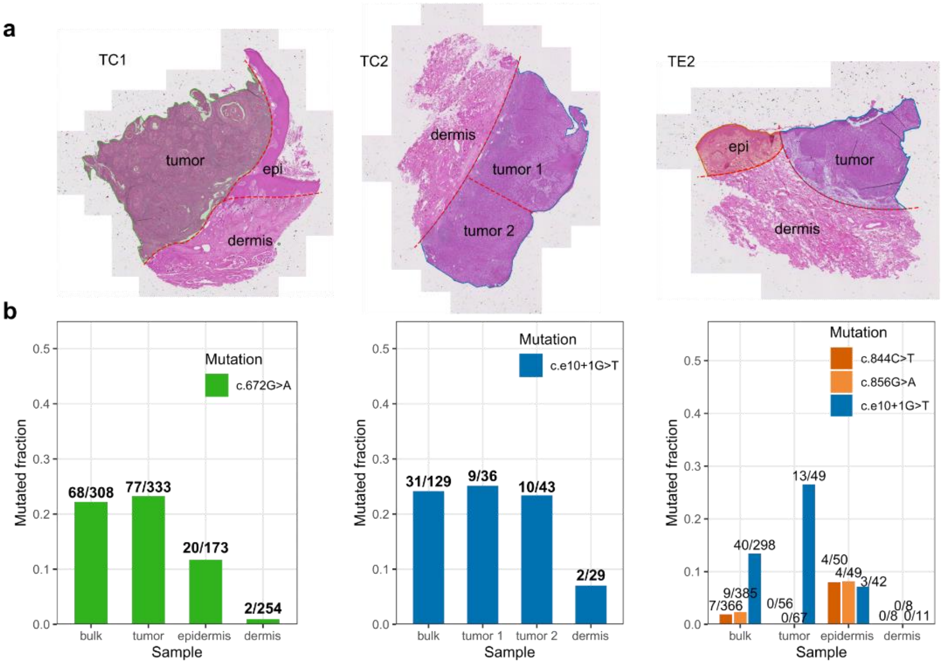


**Supplementary Fig. 7 Copy-number profiles of spatially profiled cSCC samples**

Copy-number (CN) profiles for cSCC samples. **a** CN profile across all autosomes based on whole-exome sequencing as determined by *ASCAT*. *TP53* is mapped to its chromosomal location on the x-axis. **b** Chromosome 17 copy number spot profiles in the spatially profiles cSCC samples. *TP53* mapped to its chromosomal location (left). Visualization of the average chromosome 17 copy number signal, split according to UMAP cluster defined from Fig. 5b (right). Copy-numbers profiled using *InferCNV*.

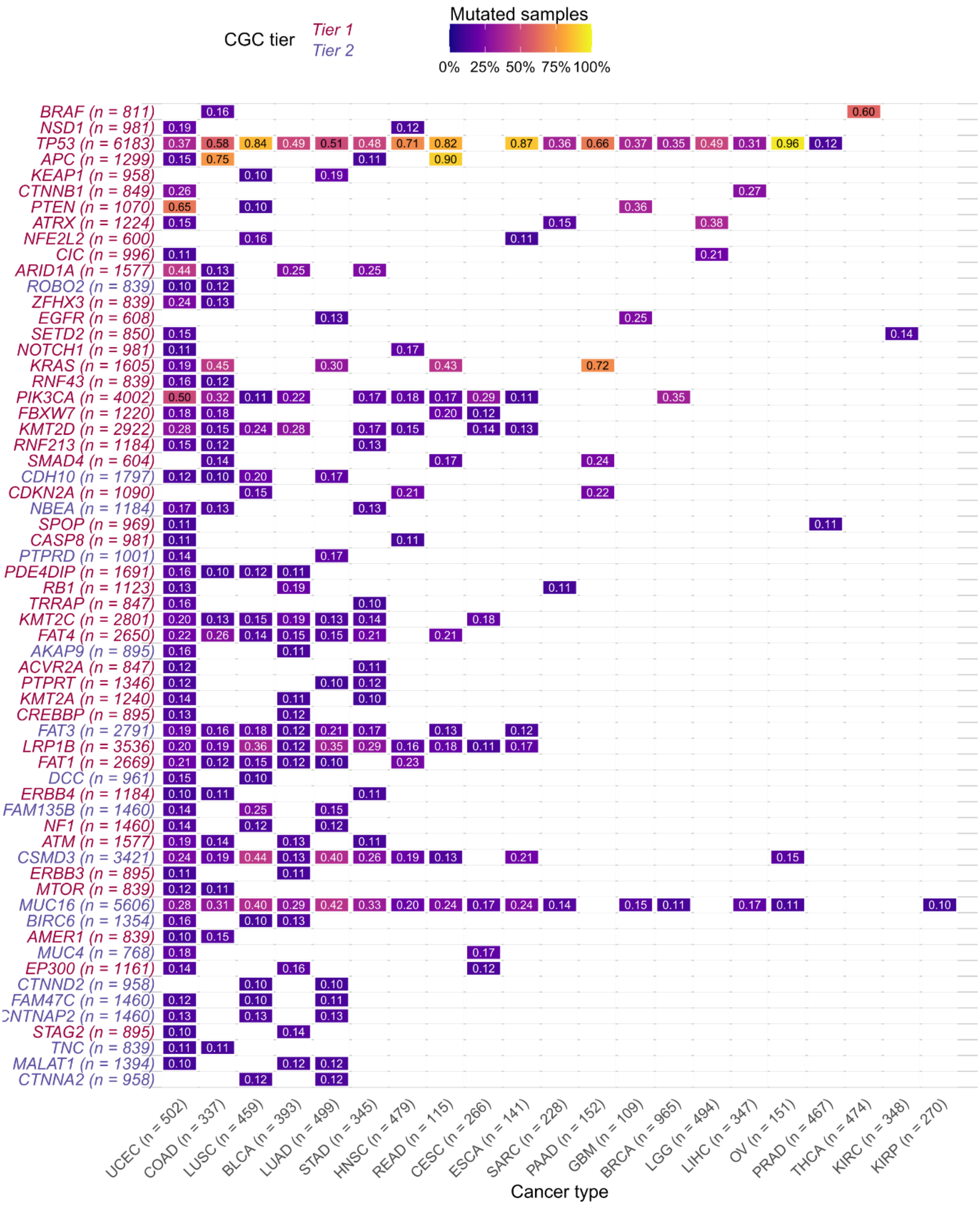


**Supplementary Fig. 8 TCGA cancer-types used to train SPOT-Mut**

Heatmap showing the TCGA cancer types used to train SPOT-Mut for each of the 62 CGC driver genes. ‘n’ equals the total number of samples for each gene or cancer type. The heatmap values show the mutation frequency for the respective gene. Gene CGC tier is indicated.

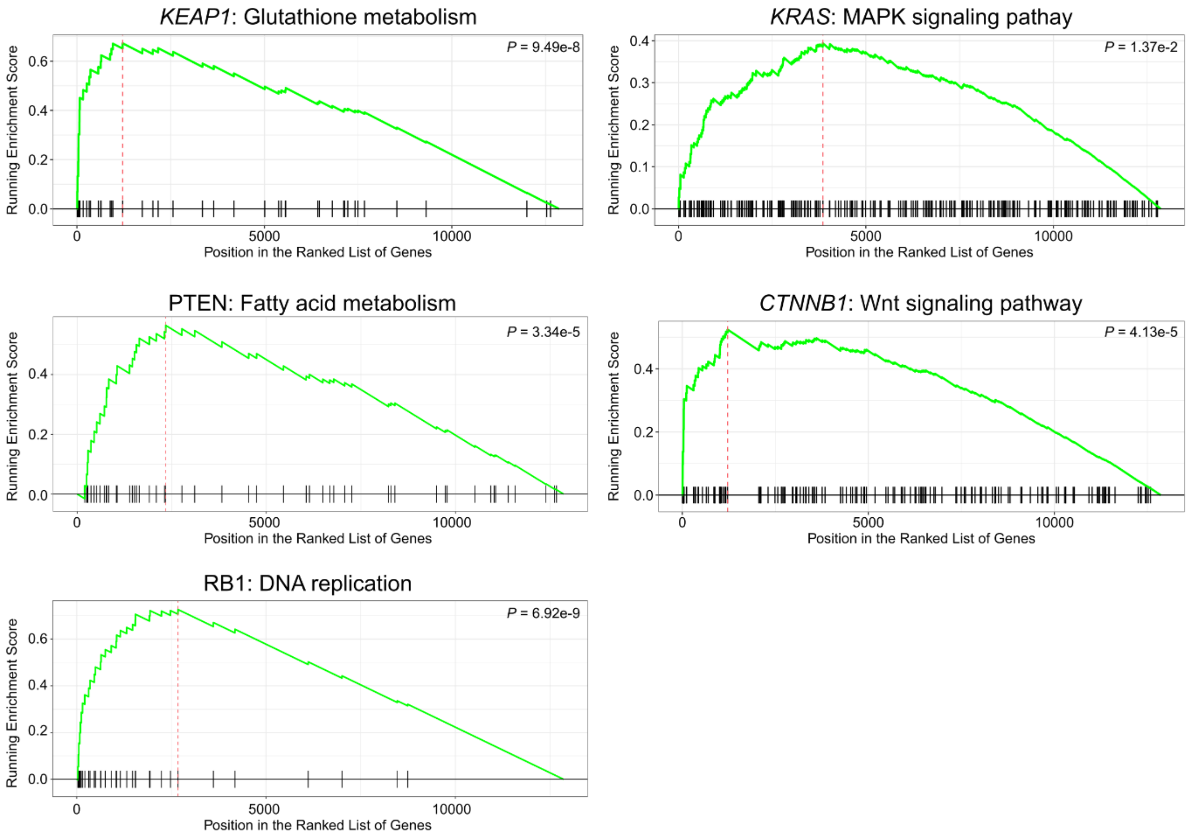


**Supplementary Fig. 9 Running score plots for *KEAP1* and *KRAS***

Gene set enrichment running score plots for KEGG gene sets biologically relevant for *KEAP1*, *KRAS*, *APC* and *CTNNB1* driver genes. Gene ranking scores were obtained from the guided backpropagation attribution scores.

### SUPPLEMENTARY TABLES

#### Supplementary Table 1: TCGA cancer type inclusion overview

Overview of the different TCGA cancer types used for developing SPOT-Mut, along with class balance and SPOT-Mut performance metrics.

#### Supplementary Table 2: WES mutation table

Mutation table listing all mutations found by Mutect2 in the bulk WES data of TC1, TC2 and TE2.

#### Supplementary Table 3: CGC driver gene screening metrics

Performance and dataset metrics for included CGC driver genes for the screening. Performance is reported as AUROC and AUPRC values evaluated for TCGA data using a GLM and for ST simulations based on both TCGA and CCLE data. For the GLM, balanced accuracy (BA) values are reported.
